# Lysine deficiency within a conserved lysine desert is critical for EEL-1/HUWE1 to support ubiquitin proteasome system function

**DOI:** 10.64898/2025.12.29.696916

**Authors:** Katherine S. Yanagi, Brenda J. Chen, Sheikh Omar Kunjo, Irini Topalidou, Nicolas Lehrbach

## Abstract

The ubiquitin proteasome system (UPS) is the primary mechanism for targeted protein degradation in eukaryotic cells. Dysfunction of this system is a driver of human disease and a hallmark of aging and late-onset neurodegenerative disorders. Understanding the mechanisms that ensure robust protein turnover may provide new avenues for treatment in these contexts. E3 ubiquitin ligases play critical roles in supplying ubiquitinated substrates to the proteasome, with HUWE1 being an enormous, versatile, and highly conserved member of this family. Here, we show that the *C. elegans* HUWE1 ortholog, EEL-1, contributes to robust protein turnover during challenges to the proteolytic capacity of the proteasome. We demonstrate that the ability of EEL-1/HUWE1 to safeguard protein turnover requires ubiquitin-binding domains within the substrate-binding arena and the HECT-type ubiquitin ligase activity, supporting a model in which EEL-1 ensures degradation by increasing ubiquitination of pre-ubiquitinated substrates. EEL-1 contains extensive lysine-deficient regions, found at conserved locations in its substrate-binding arena. Through unbiased mutagenesis screening and precise engineering of the EEL-1 protein, we uncover that introducing lysine residues into these regions is detrimental to UPS function. Together, our findings indicate a central and evolutionarily ancient role for EEL-1/HUWE1 in maintaining optimal UPS function and support targeting this E3 for therapeutic manipulation.

## INTRODUCTION

Rapid and selective protein turnover is needed for cells to maintain protein quality control via removal of damaged and misfolded proteins, and to remodel the proteome in response to environmental changes or developmental cues. Most selective protein degradation is carried out by the ubiquitin proteasome system (UPS). In eukaryotes, the canonical features of the UPS are highly conserved. Proteins are marked for degradation by post-translational conjugation of the small protein ubiquitin (Ub) to lysine residues via isopeptide linkages, a process facilitated by ubiquitin ligases [1]. Ub itself undergoes ubiquitination leading to the synthesis of substrate-anchored polyubiquitin chains [2]. Polyubiquitinated proteins are recruited to the proteasome, an elaborately regulated 33-subunit protease, which is responsible for their proteolytic destruction [3]. The UPS is precisely regulated to ensure cellular protein homeostasis is maintained across differing cell types and to allow dynamic adaptation during developmental transitions or following environmental challenges. However, the regulatory logic by which this is achieved is not fully understood [4]. Misregulation of the UPS is a common feature of disease and a hallmark of aging [5, 6]. As such, improved understanding of the UPS regulatory network may lead to new strategies for disease treatment or prevention [7–9].

The SKN-1A/Nrf1 transcription factor is the master regulator of proteasome biogenesis in animal cells and an essential component of the UPS regulatory network [10]. SKN-1A/Nrf1 is required for adequate proteasome subunit gene expression and is able to boost proteasome subunit biogenesis, allowing cells to compensate for impaired proteasome function or an increased burden of protein misfolding [11–14].

Deglycosylation of SKN-1A/Nrf1 by PNG-1/NGLY1 (a cytosolic peptide:N-glycanase enzyme) is essential for the function of the SKN-1A/Nrf1 pathway [13–15]. In *C. elegans* animals lacking either SKN-1A or PNG-1, proteasome levels are reduced, causing sensitivity to proteotoxic stress, age-dependent defects in tissue homeostasis, and reduced lifespan [14–19]. In humans, the rare genetic disease NGLY1 deficiency causes inactivation of Nrf1, and a range of symptoms associated with proteasome dysfunction [20–23].

EEL-1/HUWE1/Tom1 is an evolutionarily conserved HECT-domain E3 ubiquitin ligase, involved in the degradation of many proteins linked to a wide array of cellular, developmental, and physiological processes [24–45]. The considerable functional versatility of this ligase may be achieved through modular substrate selection; the C-terminal HECT-domain is adjacent to a giant ring-shaped arena that is decorated by several substrate-binding modules, allowing HUWE1 to select diverse substrate proteins for ubiquitination [46–48]. HUWE1 is often mutated and/or overexpressed in cancers and many HUWE1 targets are functionally implicated in tumorigenesis [49].

Further, mutations in HUWE1 cause a severe neurodevelopmental syndrome [50–52] and HUWE1 has been implicated in clearance of protein aggregates and aggregation-prone proteins associated with neurodegenerative diseases [25, 26, 45]. These connections to various diseases make HUWE1 a promising candidate for therapeutic manipulation, but a deeper understanding of how HUWE1 functions are orchestrated *in vivo* is still needed.

In a mutagenesis screen for genetic interactors of the Nrf1 pathway, we identified numerous alleles affecting *eel-1*, the *C. elegans* HUWE1/Tom1 ortholog. This finding suggests that EEL-1/HUWE1-mediated ubiquitination and SKN-1A/Nrf1-dependent transcriptional control of proteasome levels are independent yet complementary mechanisms that promote optimal UPS function. Consistently, we show that HUWE1/EEL-1 promotes degradation of a model proteasome substrate when proteasomal proteolysis is impaired. We find that the ubiquitin-binding modules and HECT ubiquitin ligase domains of EEL-1 are required for its function, supporting a model in which its ubiquitin-dependent ubiquitin ligase activity promotes optimal protein turnover via ubiquitin chain extension and/or amplification. Interestingly, EEL-1/HUWE1, like many proteins connected to UPS function, contains enigmatic ‘lysine deserts’-extended regions that are entirely devoid of lysine residues. Here, we demonstrate that the absence of lysine within these regions is critical for EEL-1 function. Collectively, our data reveal EEL-1/HUWE1 as a key component within the UPS regulatory network that enables optimal protein turnover. This work suggests that manipulation of HUWE1 activity or specificity could be widely applied to ameliorate UPS dysfunction in disease.

## RESULTS

### EEL-1/HUWE1 promotes degradation of ubiquitin-fused GFP

Reporter proteins fused at their N-terminus to the G76V mutant form of Ub are subject to rapid degradation via the Ub fusion degradation (UFD) pathway and provide a powerful tool to study proteasome-mediated degradation *in vivo* [53–57]. UFD depends on multiple Ub ligases that act in concert to extend Ub chains on the fused ubiquitin moiety [56, 58–63]. UFD is remarkably robust to inhibition of the proteolytic capacity of the proteasome; there is no detectable accumulation of Ub[G76V]-fused GFP in cancer cells unless >80% of proteasome activity is eliminated [53]. These observations suggest that the UPS is highly robust to proteolytic challenges. However, the mechanistic basis and physiologic significance of robust UPS function in this context is not clear.

We previously generated a transgene that ubiquitously expresses Ub[G76V]::GFP (hereafter UbV-GFP) in *C. elegans* [14, 16]. UbV-GFP fluorescence is not detectable in wild type animals but accumulates to low but detectible levels in intestinal cells of *skn-1a* and *png-1* mutant animals, reflecting a requirement for SKN-1A/Nrf1 for adequate proteasome levels [14]. We hypothesized that UbV-GFP accumulation in SKN-1A pathway mutants would be exacerbated if other components that promote robust UPS function are inactivated. To uncover these factors, we carried out a forward genetic screen for increased UbV-GFP fluorescence in animals lacking either *skn-1a* or *png-1*. This screen identified several alleles affecting the *eel-1* gene (Fig 1A). *eel-1* encodes a highly conserved ∼465 kDa HECT-domain E3 ubiquitin ligase orthologous to human HUWE1 and *S. cerevisiae* Tom1. HUWE1 knockdown in human cancer cells causes increased accumulation of UbV-GFP [58] and inactivation of Tom1 in yeast causes partial stabilization of UFD substrates [60]. RNAi-mediated knockdown of *eel-1* leads to increased UbV-GFP accumulation in *C. elegans* using an independently generated UbV-GFP transgene [55]. Thus, identification of *eel-1* in our screen supports that this ubiquitin ligase plays a conserved role in degradation of Ub fusion proteins.

**Figure 1.**
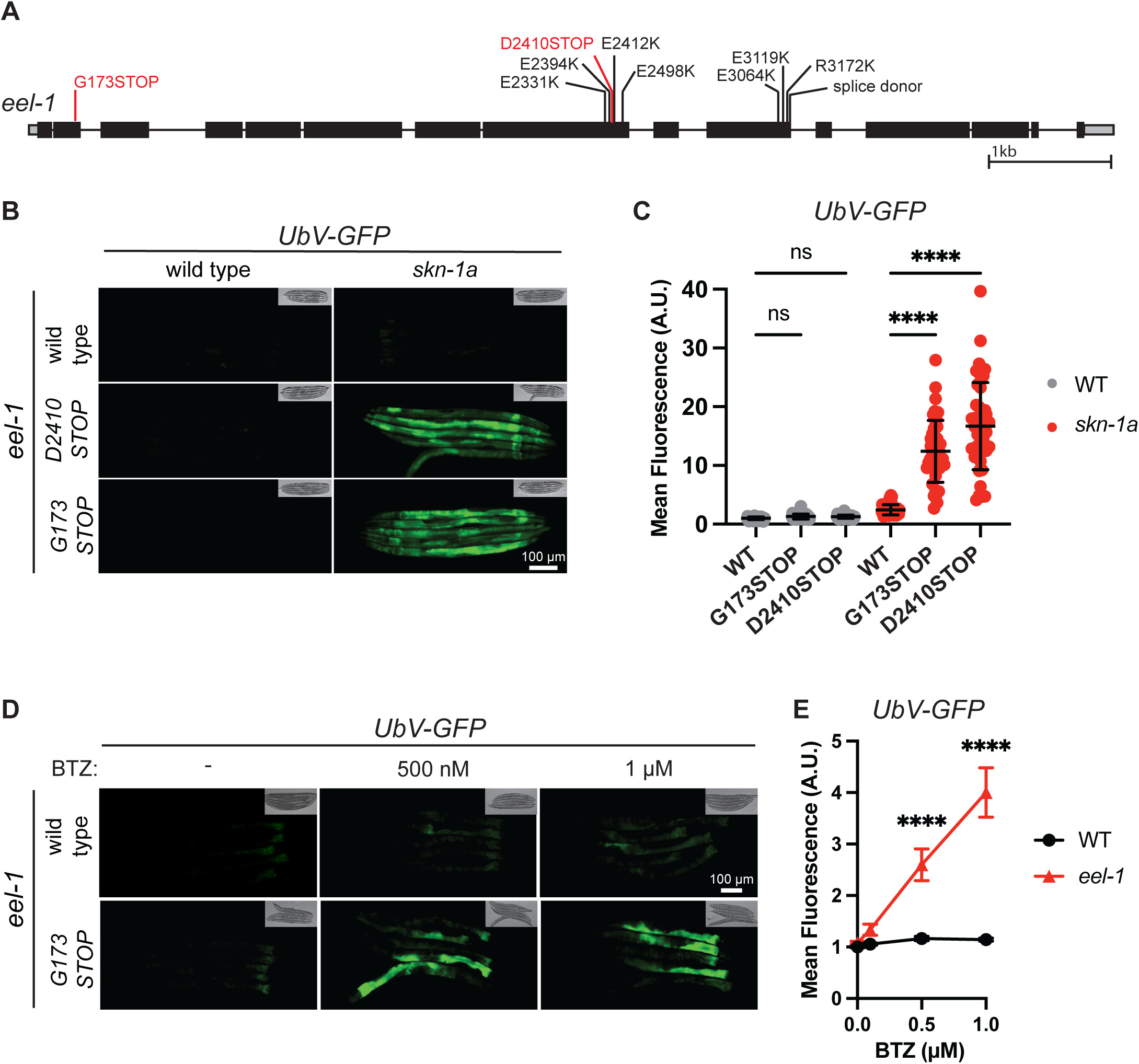
EEL-1/HUWE1 promotes turnover of UbV-GFP A) Schematic of the *eel-1* gene. Locations and effects of mutations identified in a mutagenesis screen for increased accumulation of UbV-GFP are labelled in black. Locations and effects of CRISPR/Cas9-engineered alleles are labelled in red. B) Fluorescence micrographs showing that simultaneous inactivation of *skn-1a* and *eel-1* causes dramatic accumulation of UbV-GFP. All images are of L4 stage animals. Scale bar 100 μM. C) Quantification of UbV-GFP levels shown in (B). UbV-GFP accumulation is not increased in *eel-1* single mutants and is moderately increased in *skn-1a* single mutants. UbV-GFP accumulation is dramatically increased in *eel-1 skn-1a* double mutants. Error bars show mean ± SD. n=45 animals per genotype. Ns p>0.05, **** p<0.0001, ordinary one-way ANOVA with Šídák’s multiple comparisons test. D) Fluorescence micrographs showing that accumulation of UbV-GFP following proteasome inhibition is increased in *eel-1* mutants. L4 animals were exposed to the indicated BTZ concentration (or DMSO control) for 24 hours prior to imaging. Imaged animals are day 1 adults. Scale bar 100 μM. E) Quantification of UbV-GFP levels following BTZ exposure as described in panel (D). Exposure of *eel-1* mutant animals to 0.5-1 μM BTZ leads to increased accumulation of UbV-GFP but has no effect on the wild type. Error bars show mean ± SD. N=30 animals per genotype and condition (BTZ concentrations tested were: DMSO control, 0.1 μM, 0.5 μM, 1 μM). P-values indicate comparison between *eel-1* mutant and wild type for each drug concentration. Ns p>0.05, **** p<0.0001, ordinary two-way ANOVA with Šídák’s multiple comparisons test.

To validate our screening results, we used CRISPR/Cas9 to introduce a premature termination codon/frameshift cassette at two positions in the *eel-1* coding sequence (G173STOP, D2410STOP; Fig 1A). Strikingly, in a wild type genetic background, neither allele causes any accumulation of UbV-GFP. However, in a *skn-1a* or *png-1* mutant background, *eel-1* inactivation causes very high levels of UbV-GFP, far exceeding levels observed in single mutants (Fig 1B-C; Fig S1). We conclude that EEL-1 promotes proteasomal degradation of UbV-GFP but is not essential for this process. Rather, the contribution of EEL-1 becomes salient when SKN-1A-mediated control of proteasome levels is lost.

This genetic interaction suggested that EEL-1 could play an important role to ensure turnover of ubiquitin fusion proteins when proteasome levels or function is reduced. The proteasome inhibitor bortezomib (BTZ) impairs proteasome function by blocking the β5/chymotrypsin-like active site, which is rate-limiting for protein turnover [64, 65]. In WT animals, exposure to BTZ at high concentrations causes accumulation of UbV-GFP [16]. We found that in animals lacking EEL-1, the concentration of BTZ required to cause accumulation of UbV-GFP is significantly lower compared to the wild type (Fig 1D-E). We conclude that EEL-1 is important to ensure turnover of UbV-GFP in contexts where the capacity for proteasome-mediated protein turnover is impaired.

### Simultaneous inactivation of EEL-1 and SKN-1A causes severe defects in development and physiology

Inactivation of EEL-1 causes a range of developmental and physiological defects [39–42]. During our analysis, we noticed that adverse consequences of *eel-1* inactivation are severely exacerbated by loss of the SKN-1A/Nrf1 pathway. In *eel-1 skn-1a* or *png-1*; *eel-1* double mutants, we observed frequent vulval rupture during the first two days of adulthood (Fig 2A, B). Additional rupture events occur later and almost all double mutant animals rupture by day 4 (Fig 2C). In contrast, single mutants lacking *eel-1* rarely rupture within the first two days of adulthood, but do show rupture later in adulthood, affecting ∼30% of animals by day 4 (Fig 2A-C). *skn-1a* and *png-1* mutants show elevated vulval rupture later in life (by day 7) [16], but no rupture during the first 4 days of adulthood (Fig 2A-C). Thus, EEL-1 and SKN-1A each contribute to maintenance of vulval integrity during adulthood and simultaneous inactivation of both is synergistically detrimental. In most *png-1; eel-1* mutant animals, the vulva ruptures prior to significant progeny production and the progeny produced rarely develop into viable adults. As a result, we could not maintain a double mutant population for more than one generation and have not further characterized genetic interactions between *png-1* and *eel-1*.

**Figure 2.**
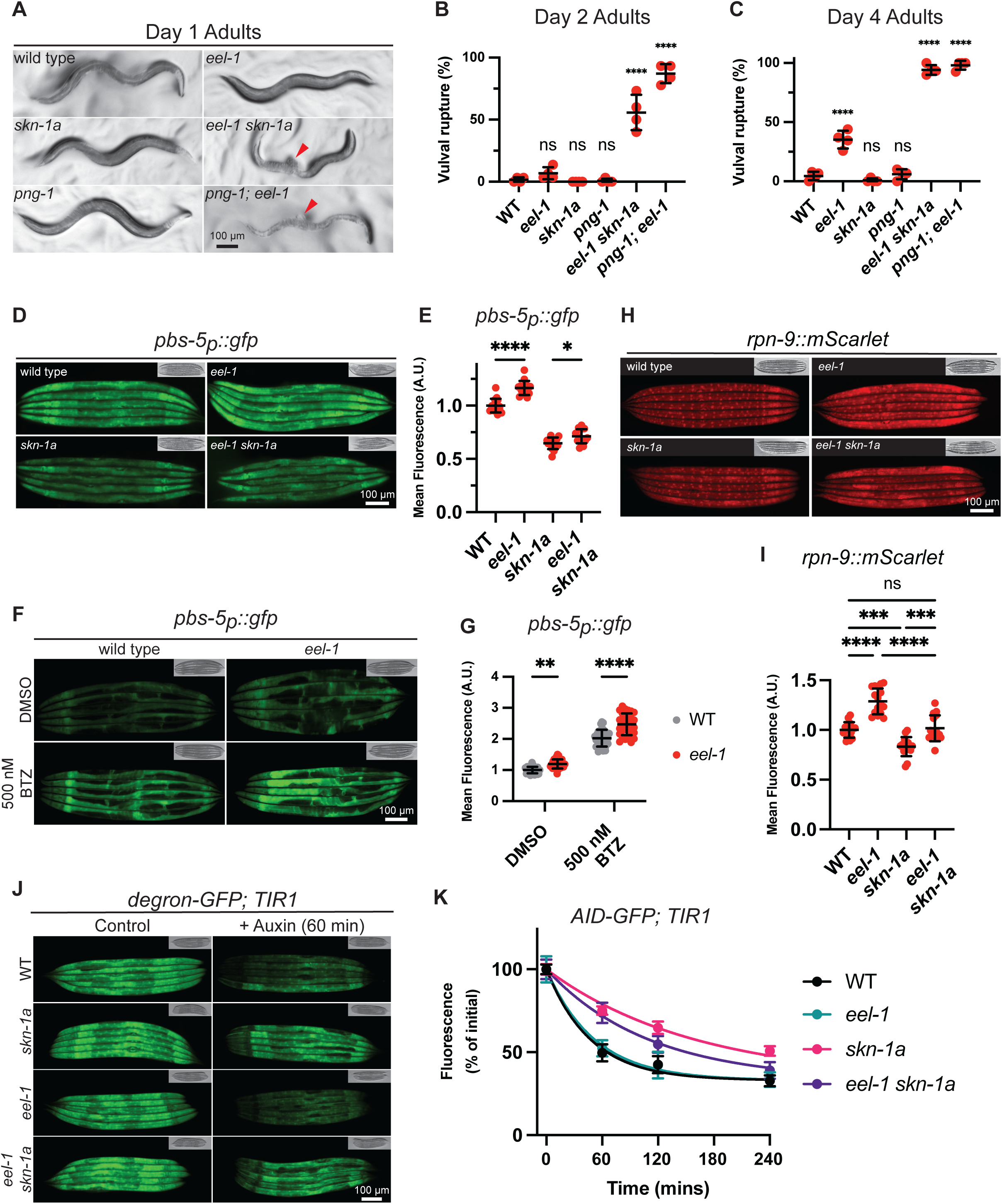
Simultaneous inactivation of EEL-1/HUWE1 and the SKN-1A/Nrf1 pathway causes severe defects in development and physiology that are not caused by reduced proteasome levels. A) Micrographs showing day 1 adult animals. Red arrows indicate vulval rupture. Vulval rupture occurs frequently in *eel-1 skn-1a* and *png-1*; *eel-1* double mutants but is not observed in wild type animals or in single mutants. Scale bar 100 μM. B) Quantification of vulval rupture in day 2 adults. Results are shown for n=4 replicate experiments, >25 animals were assayed in each replicate experiment. Error bars show mean ± SD. Ns p>0.05, **** p<0.0001, indicates comparison to the WT control, ordinary one-way ANOVA with Šídák’s multiple comparisons test. C) Quantification of vulval rupture in day 4 adults. Results are shown for n=4 replicate experiments, >25 animals were assayed in each replicate experiment. Error bars show mean ± SD. Ns p>0.05, **** p<0.0001, indicates comparison to the WT control, ordinary one-way ANOVA with Šídák’s multiple comparisons test. D) Fluorescence micrographs showing expression of the *pbs-5_p_::gfp* transcriptional reporter of proteasome subunit gene expression. Expression of the reporter is increased in *eel-1* mutant animals and reduced in *skn-1a* mutant animals. Expression of the reported is not further reduced in *eel-1 skn-1a* double mutants. Images show L4 stage animals. Scale bar 100 μM. E) Quantification of *pbs-5_p_::gfp* reporter expression shown in (D). Expression of the reporter is increased in *eel-1* mutant animals compared to the wild type. Expression is reduced in *skn-1a* mutants. Although expression in *eel-1 skn-1a* double mutants is reduced compared to wild type, it is slightly increased compared to *skn-1a* single mutants. Results are shown for n=15 animals. Error bars show mean ± SD. * p<0.05, **** p<0.0001, ordinary one-way ANOVA with Tukey’s multiple comparisons test. F) Fluorescence micrographs showing expression of the *pbs-5_p_::gfp* transcriptional reporter of proteasome subunit gene expression following exposure to BTZ. L4 animals were exposed to 500 nM BTZ (or DMSO control) for 24 hours prior to imaging. Imaged animals are young adults. Expression of the *pbs-5_p_::gfp* reporter is increased in *eel-1* mutants as compared to the wild type under both control (DMSO) conditions and following BTZ exposure. Scale bar 100 μM. G) Quantification of *pbs-5_p_::gfp* reporter expression shown in (F). Expression of the *pbs-5_p_::gfp* GFP reporter is higher in *eel-1* mutants in control conditions and following BTZ exposure. Results are shown for n=30 animals for each genotype/condition. Error bars show mean ± SD. ** p<0.01, **** p<0.0001, ordinary two-way ANOVA, uncorrected Fisher’s LSD. H) Fluorescence micrographs showing levels of an endogenously tagged proteasome subunit, RPN-9, tagged with mScarlet. Proteasome levels are increased in *eel-1* mutants and reduced in *skn-1a* mutants, but are not further reduced in *eel-1 skn-1a* double mutants. Images show L4 stage animals. Scale bar 100 μM. I) Quantification of levels of the RPN-9 proteasome subunit shown in (H). RPN-9 levels are increased in *eel-1* mutants and reduced in *skn-1a* mutants. RPN-9 levels are unchanged compared to the wild type in *eel-1 skn-1a* double mutants. Results are shown for n=15 animals per genotype. Error bars show mean ± SD. Ns p>0.05, *** p<0.001, **** p<0.0001, ordinary one-way ANOVA with Šídák’s multiple comparisons test. J) Fluorescence micrographs showing auxin-induced degradation of degron-GFP. Images show L4 stage animals either before, or 60 minutes after, transfer to plates supplemented with 50 mM auxin. Auxin-induced degradation of degron-GFP is reduced in *skn-1a* mutants but not in *eel-1* mutants. K) Quantification of degron-GFP fluorescence following auxin exposure under the same conditions shown in (J); animals were imaged over a 4-hour period following auxin exposure. Destruction of degron-GFP is delayed in *skn-1a* mutants but is not affected by *eel-1*. For each genotype, fluorescence intensity is normalized to the initial (pre-auxin-exposure) value to calculate the percent fluorescence remaining at each time point. N=40-45 animals per genotype at each time-point. Error bars show mean ± SD. Trend line shows non-linear regression (one phase decay). degron-GFP half-life in each genotype (95% confidence interval, mins): WT: 28-38; *skn-1a*: 101-121; *eel-1*: 28-44; *eel-1 skn-1a*: 66-88. In all panels of this figure, *eel-*1 denotes the *eel-1*[*G173STOP*] mutant.

We also noticed that *eel-1* mutants show a marked delay in larval growth and development, a defect that is particularly severe at elevated temperatures (Fig S2). *skn-1a* single mutants develop at the same rate as the wild type, but the growth delay caused by *eel-1* inactivation is strongly enhanced in double mutants (Fig S2). These data indicate that *eel-1* is required for timely developmental progression, and *skn-1a* becomes necessary to support development when *eel-1* is inactive. We conclude that abnormalities in development and physiology caused by inactivation of the EEL-1 ubiquitin ligase are exacerbated by loss of proteasome regulation by SKN-1A. Given the synergistic impact of *eel-1* and *skn-1a* mutations on UbV-GFP levels, these data suggest that simultaneous loss of EEL-1/HUWE1 and SKN-1A/Nrf1 severely impairs the UPS, causing major developmental and physiological defects.

### EEL-1 is not required to maintain adequate proteasome levels

The strong genetic interaction between *eel-1* and *skn-1a* suggests that SKN-1A and EEL-1 could act in separate pathways that independently enhance proteasome levels or activity. To examine whether EEL-1 impacts transcriptional control of proteasome levels, we generated a reporter in which the promoter of the *pbs-5* gene drives GFP expression (*pbs-5_p_::gfp)*. The *pbs-5* gene encodes the β5 subunit of the proteasome. PBS-5/β5 is essential for proteasome assembly and function, is regulated by SKN-1A, and is likely rate-limiting for proteasome function *in vivo* [14, 17, 66–69]. We find that loss of *eel-1* does not reduce, but rather increases, *pbs-5_p_::gfp* expression and does not further decrease *pbs-5 _p_::gfp* expression in *skn-1a* mutants (Fig 2D-E). Exposure to BTZ leads to increased *pbs-5_p_::gfp* expression in wild type animals, as expected, and the extent of activation is higher in *eel-1* mutants (Fig 2F-G). Therefore, the dramatic UbV-GFP accumulation in *eel-1 skn-1a* double mutants is not explained by reduced expression of proteasome subunit genes.

EEL-1 could impact proteasome levels via a post-transcriptional effect not captured by the *pbs-5_p_::gfp* reporter. We therefore examined animals in which an endogenous proteasome subunit is tagged with mScarlet (*rpn-9::mScarlet*) [70]. Supporting observations made with the *pbs-5_p_::gfp* transcriptional reporter for proteasome subunit gene expression, mScarlet-tagged RPN-9 protein levels are increased in *eel-1* mutant animals compared to the wild type (Fig 2H, I). As expected, *skn-1a* mutants show reduced RPN-9 levels [14], but levels are not further reduced in the double mutants lacking both EEL-1 and SKN-1A (Fig 2H, I). Thus, the strong genetic interaction between *eel-1* and *skn-1a* is not explained by a change in proteasome abundance.

### EEL-1 is not generally required for protein degradation by the proteasome

We next sought to address the possibility that proteasome function is generally impaired in animals lacking EEL-1. To do this, we examined auxin-induced degradation (AID) of GFP. AID is a heterologous protein degradation system derived from plant hormone signaling and has been used to drive conditional proteasome-dependent protein depletion [71, 72]. GFP fused to an auxin-sensitive degron (degron-GFP; expressed in all tissues) is degraded by the proteasome following addition of auxin, triggering interaction with the ubiquitin ligase TIR1. In this system, exposure to 50 mM auxin leads to rapid destruction of degron-GFP, with diminished fluorescence readily detectable after 1 hour [73]. Consistent with reduced proteasome levels causing a general impairment of protein turnover, clearance of degron-GFP is delayed in *skn-1a* mutant animals, with a roughly 3-fold increase in degron-GFP half-life (Fig 2J, K). In contrast, auxin-induced degradation is unchanged in *eel-1* mutants. Additionally, there is no further delay in degron-GFP turnover in *eel-1 skn-1a* double mutants compared to *skn-1a* (Fig 2J, K). We conclude that EEL-1 is not required for AID of GFP. Since AID is proteasome-dependent, this argues that the strong genetic interaction between *eel-1* and *skn-1a* is not explained by a general reduction in all proteasome functions.

### EEL-1 ubiquitin ligase activity and ubiquitin-binding domains promote turnover of UbV-GFP and vulval integrity

We sought to understand the mechanism by which EEL-1 promotes degradation of UbV-GFP. EEL-1/HUWE1 contains a conserved C-terminal HECT domain responsible for transfer of ubiquitin to substrates. The ubiquitination reaction requires a conserved cysteine residue found within the HECT domain [74]. We engineered inactivating mutations in the EEL-1 HECT domain by CRISPR/Cas9 (C4144A and C4144S) (Fig 3A). These mutations dramatically enhance UbV-GFP accumulation in *skn-1a* mutants (Fig 3B, C). We therefore conclude that the E3 ubiquitin ligase activity of EEL-1 is required to promote UbV-GFP turnover, consistent with a conserved role of EEL-1/HUWE1/Tom1 as a ubiquitin ligase in the UFD pathway [58, 60].

**Figure 3.**
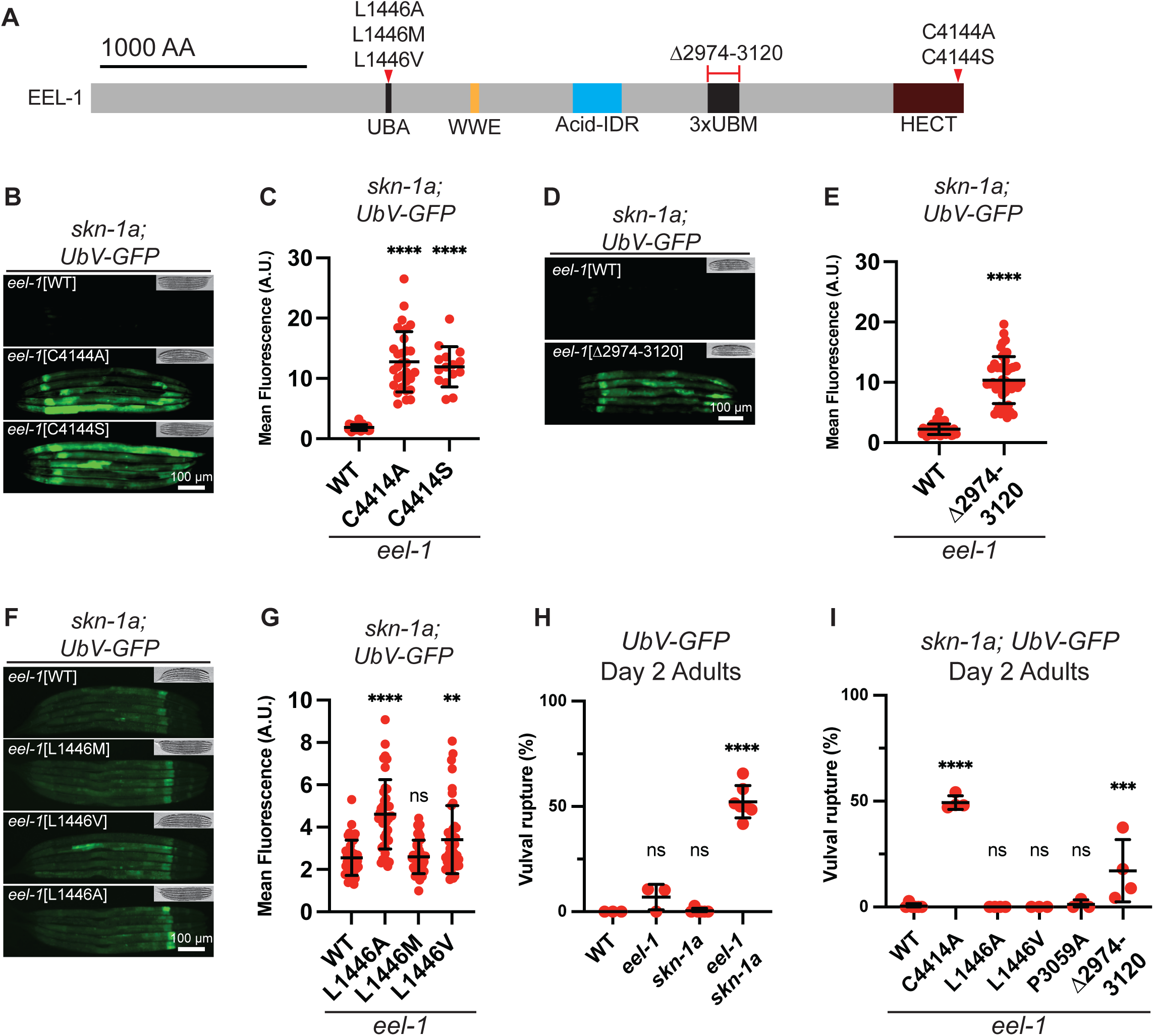
EEL-1/HUWE1 ubiquitin ligase activity and ubiquitin-binding domains are required to promote turnover of UbV-GFP and vulval integrity. A) Schematic of the EEL-1 protein showing domain architecture and locations of mutations studied in this figure. B) Fluorescence micrographs showing that mutation of the EEL-1 active site causes accumulation of UbV-GFP in a *skn-1a* mutant background. Images show L4 stage animals. Scale bar 100 μM. C) Quantification of UbV-GFP levels shown in (B). All animals are *skn-1a* mutant, the x-axis label indicates the *eel-1* genotype. Results are shown for nζ15 animals per genotype. Error bars show mean ± SD. **** p<0.0001, indicates P-value for comparison to the *eel-1*[WT] control, ordinary one-way ANOVA with Dunnett’s multiple comparisons test. D) Fluorescence micrographs showing that deletion of the EEL-1 3xUBM module causes accumulation of UbV-GFP in a *skn-1a* mutant background. Scale bar 100 μM. E) Quantification of UbV-GFP levels shown in (D). All animals are *skn-1a* mutant, the x-axis label indicates the *eel-1* genotype. Results are shown for n=45 animals per genotype. Bars show mean ± SD. **** p<0.0001, indicates P-value for comparison to the *eel-1*[WT] control, Welch’s two-tailed t test. F) Fluorescence micrographs showing the effects of mutation of the EEL-1 UBA domain on UbV-GFP accumulation in a *skn-1a* mutant background. The L1446M mutation is a conservative substitution that humanizes EEL-1 at this residue. The L1446V mutation is equivalent to a patient allele and the L1446A mutation is expected to disrupt ubiquitin binding similarly to L1446V. UbV-GFP levels are slightly increased in L1446V and L1446A mutant animals. Scale bar 100 μM. G) Quantification of UbV-GFP levels shown in (F). All animals are *skn-1a* mutant, the x-axis label indicates the *eel-1* genotype. Results are shown for n=45 animals per genotype. Error bars show mean ± SD. Ns p>0.5, ** p<0.01, **** p<0.0001, indicates P-value for comparison to the *eel-1*[WT] control, ordinary one-way ANOVA with Dunnett’s multiple comparisons test. H) Quantification of vulval rupture phenotypes caused by *skn-1a* and *eel-1* null mutations in animals carrying the UbV-GFP transgene. The *eel-1* mutation tested is *eel-1*[*G173STOP*]. Inactivation *eel-1* and *skn-1a* has the same effect on vulval integrity in the UbV-GFP transgenic background as in non-transgenic animals (compare to Figure 2B). Results are shown for n=3-7 replicate experiments, 25-35 animals were assayed in each replicate experiment. Error bars show mean ± SD. Ns p>0.05, **** p<0.0001, indicates P-value for comparison to the *eel-1*[WT] control, ordinary one-way ANOVA with Šídák’s multiple comparisons test. I) Quantification of vulval rupture phenotypes caused by various *eel-1* mutations. All animals are *skn-1a* mutant and carry the UbV-GFP transgene, the x-axis label indicates the *eel-1* genotype. Results are shown for n=3-4 replicate experiments, 25-35 animals were assayed in each replicate experiment. Error bars show mean ± SD. Ns p>0.05, *** p<0.001, **** p<0.0001, indicates P-value for comparison to the *eel-1*[WT] control, ordinary one-way ANOVA with Šídák’s multiple comparisons test. Additional statistical analysis of these data is provided in Table S1.

EEL-1/HUWE1/Tom1 contains conserved ubiquitin-binding domains including a ubiquitin associated (UBA) domain and three repeated ubiquitin binding motifs (UBMs) (Fig 3A). Our screen isolated two alleles (E3064K, E3119K) affecting the UBMs. We noted that the E3064K mutation alters a conserved residue within EEL-1’s second UBM (Fig S3), near an LP dipeptide required for Ub binding [75, 76]. To directly test whether ubiquitin-binding is important for EEL-1 function, we first generated a CRISPR/Cas9 allele, P3059A, which is predicted to disrupt Ub-binding [75]. However, P3059A had no effect on UbV-GFP turnover in *skn-1a* mutant animals (Fig S4). This suggested that the 3xUBM module may not be required for EEL-1 function in UbV-GFP turnover, or the mutation does not fully disrupt its function. We therefore generated an in-frame deletion allele that removes all three UBMs (Fig 3A). This *eel-1*[*Δ2974-3120*] mutation strongly enhances accumulation of UbV-GFP (Fig 3D, E), suggesting that robust UbV-GFP turnover requires the 3xUBM Ub-binding domain of EEL-1.

A mutation that disrupts the UBA domain of HUWE1, M1328V, causes a neurodevelopmental syndrome [51, 77]. The equivalent residue in *C. elegans* EEL-1 is L1446; M1328/L1446 lies within the conserved binding interface through which various UBA domains interact with ubiquitin [78–80]. A conserved mode of Ub-binding involving M1328/L1146 is supported by sequence alignment and Alpha Fold structural prediction (Fig S5). We generated animals harboring the patient-equivalent mutation L1446V and another substitution L1146A, both of which are expected to disrupt Ub binding. We additionally generated a strain in which the UBA domain of *C. elegans* EEL-1 is ‘humanized’ by a L1446M substitution, which is not expected to disrupt Ub binding. The EEL-1[L1446A] and EEL-1[L1446V] alleles both cause a mild but significant increase in accumulation of UbV-GFP in *skn-1a* mutant animals, whereas there is no effect of the EEL-1[L1446M] substitution (Fig 3F, G). Thus, the UBA domain of EEL-1 contributes to robust UbV-GFP turnover, though the effect is modest compared to that of the 3xUBM domain. We conclude that Ub-binding domains of EEL-1 are involved in ensuring optimal degradation of UbV-GFP. These data are consistent with a model in which EEL-1 binds UbV-GFP via Ub-binding domains and promotes UbV-GFP degradation via ubiquitin-dependent ubiquitin ligase activity [58, 60].

We next asked whether the *eel-1* alleles affecting the ubiquitin ligase or Ub-binding domains have effects on vulval rupture. We carried out these experiments using animals harboring the UbV-GFP transgene, which we confirmed does not alter the rupture phenotypes caused by *eel-1* and/or *skn-1a* inactivation (Fig 3H; compare to Fig 2B). In a *skn-1a* mutant background, the C4144A HECT domain mutation caused vulval rupture to a similar extent as the *eel-1* null, while mutations affecting ubiquitin binding domains corresponded to their effects on UbV-GFP: only *eel-1*[*Δ2974-3120*] showed significant vulval rupture (Fig 3I). Thus, EEL-1 ensures turnover of UbV-GFP and promotes vulval integrity via similar mechanisms, suggesting that ubiquitin-dependent ubiquitin ligase activity of EEL-1/HUWE1 plays important roles in animal physiology by promoting optimal protein turnover.

### The Acid-IDR of EEL-1 is not required for degradation of UbV-GFP or vulval integrity

Our screen isolated a cluster of amino acid substitutions that do not affect ubiquitin binding domains or the HECT domain (E2331K, E2394K, E2412K, E2498K; Fig 4A). These substitutions cluster within a region of the protein that is highly enriched in acidic residues (∼50% glutamate or aspartate) and is intrinsically disordered (Fig 4A-C). We therefore term this region the Acid-IDR domain. The Acid-IDR domain is involved in recognition of orphaned ribosomal subunits and other basic proteins for ubiquitination and degradation [24, 46]. The alleles isolated in our screen each introduce a basic amino acid, lysine, into this domain, which could disrupt electrostatic interactions with substrates or regulators. To test the role of the Acid-IDR, we generated in-frame deletion alleles that remove all or part of the domain (Fig 4A). We observed no UbV-GFP accumulation following deletion of the Acid-IDR in a *skn-1a* mutant background (Fig 4D, E). Further, we measured vulval rupture and found no increase compared to control animals (Fig 4F). We conclude that the Acid-IDR of EEL-1 is not required for degradation of UbV-GFP or vulval integrity in *skn-1a* mutant animals.

**Figure 4.**
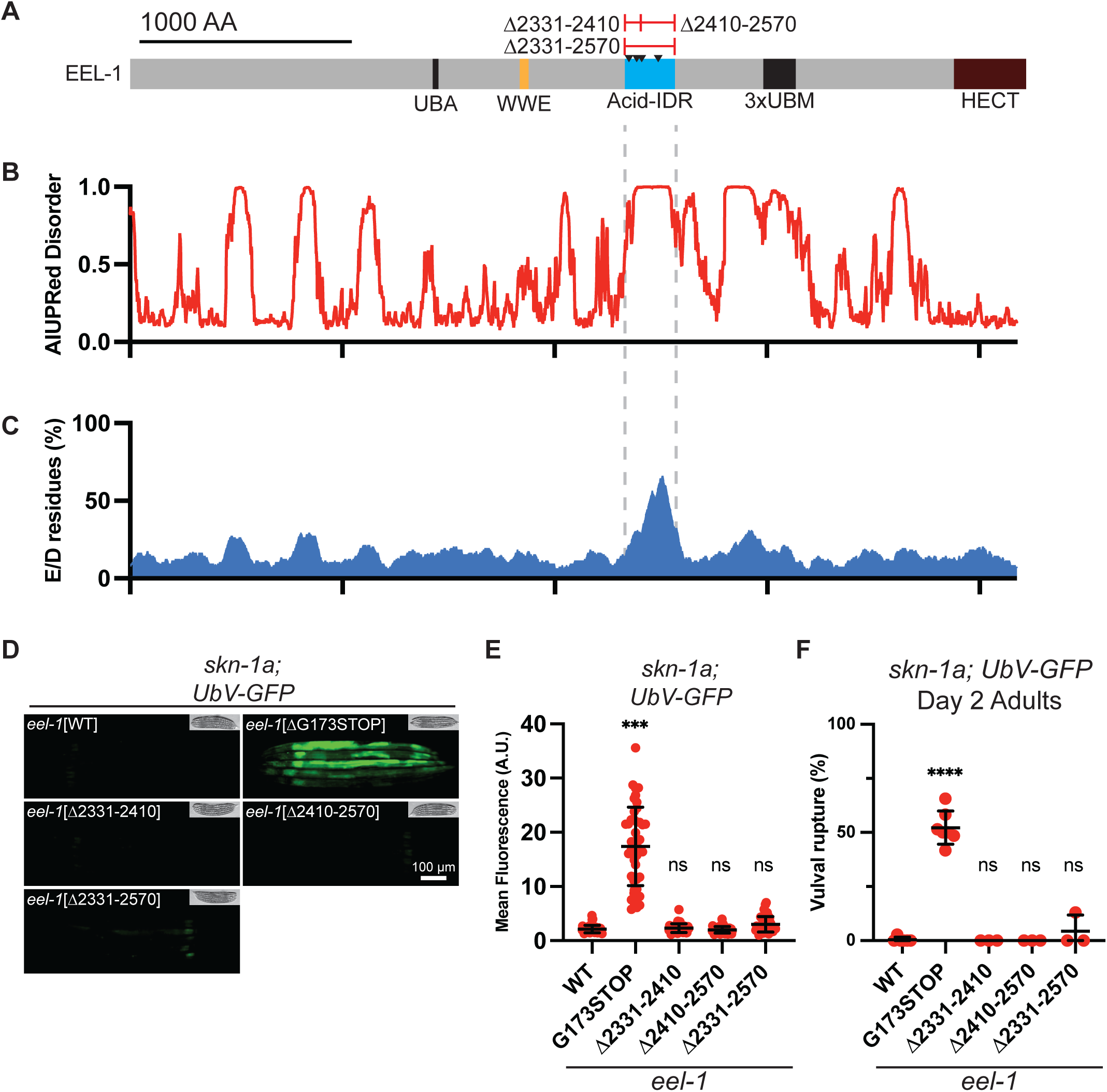
The Acid-IDR domain of EEL-1/HUWE1 is not required for UbV-GFP turnover or vulval integrity. A) Schematic of the EEL-1 protein showing domain architecture and locations of in-frame deletion mutations studied in this figure. Black arrows indicate locations of amino acid substitutions identified in our screen for increased UbV-GFP accumulation (E2331K, E2394K, E2412K, E2498K). B) The EEL-1 Acid-IDR corresponds to a region of predicted disorder as calculated by AIUPred. C) Amino acid composition of EEL-1, showing enrichment of acidic residues in the Acid-IDR. The percent acidic amino acids (glutamate or aspartate) in a 100 amino acid window around each position is plotted. D) Fluorescence micrographs showing that deletion of the Acid-IDR of EEL-1 does not lead to increased accumulation of UbV-GFP in *skn-1a* mutants. The G173STOP mutant is included as a positive control. Scale bar 100 μM. E) Quantification of UbV-GFP levels shown in (D). All animals are *skn-1a* mutants, the x-axis label indicates the *eel-1* genotype. Results are shown for n=45 animals per genotype. Error bars show mean ± SD. Ns p>0.5, *** p<0.001, indicates P-value for comparison to the *eel-1*[WT] control, ordinary one-way ANOVA with Dunnett’s multiple comparisons test. F) Quantification of vulval rupture phenotypes caused by various *eel-1* mutations affecting the Acid-IDR domain. All animals are *skn-1a* mutants and carry the UbV-GFP transgene, the x-axis label indicates the *eel-1* genotype. Results with the *eel-1*[*G173STOP*] allele are included for comparison. Results are shown for n=3-7 replicate experiments, 25-35 animals were assayed in each replicate experiment. Error bars show mean ± SD. Ns p>0.05, **** p<0.0001, indicates P-value for comparison to the *eel-1*[WT] control, ordinary one-way ANOVA with Šídák’s multiple comparisons test. Additional statistical analysis of these data is provided in Table S1.

### Lysine desert mutations in eel-1 disrupt turnover of UbV-GFP

Interestingly, our unbiased mutagenesis screen identified several amino acid substitutions that introduce lysine (K) residues in two distinct clusters (Fig 5A). The clusters are in or nearby to the Acid-IDR (E2331K, E2394K, E2412K, E2598K) and 3xUBM domains (E3064K, E3119K, R3172K). Comparing the sequence composition and domain organization of EEL-1/HUWE1/Tom1 orthologs across diverse eukaryotes, these domains are positioned within deeply conserved lysine-deficient regions (termed lysine deserts; Fig 5A, S6, S7). Lysine deserts are frequently found in UPS components and prevent their inactivation via accidental ubiquitination [81–86]. We therefore hypothesized that the absence of lysine from these domains is required for EEL-1 function.

**Figure 5.**
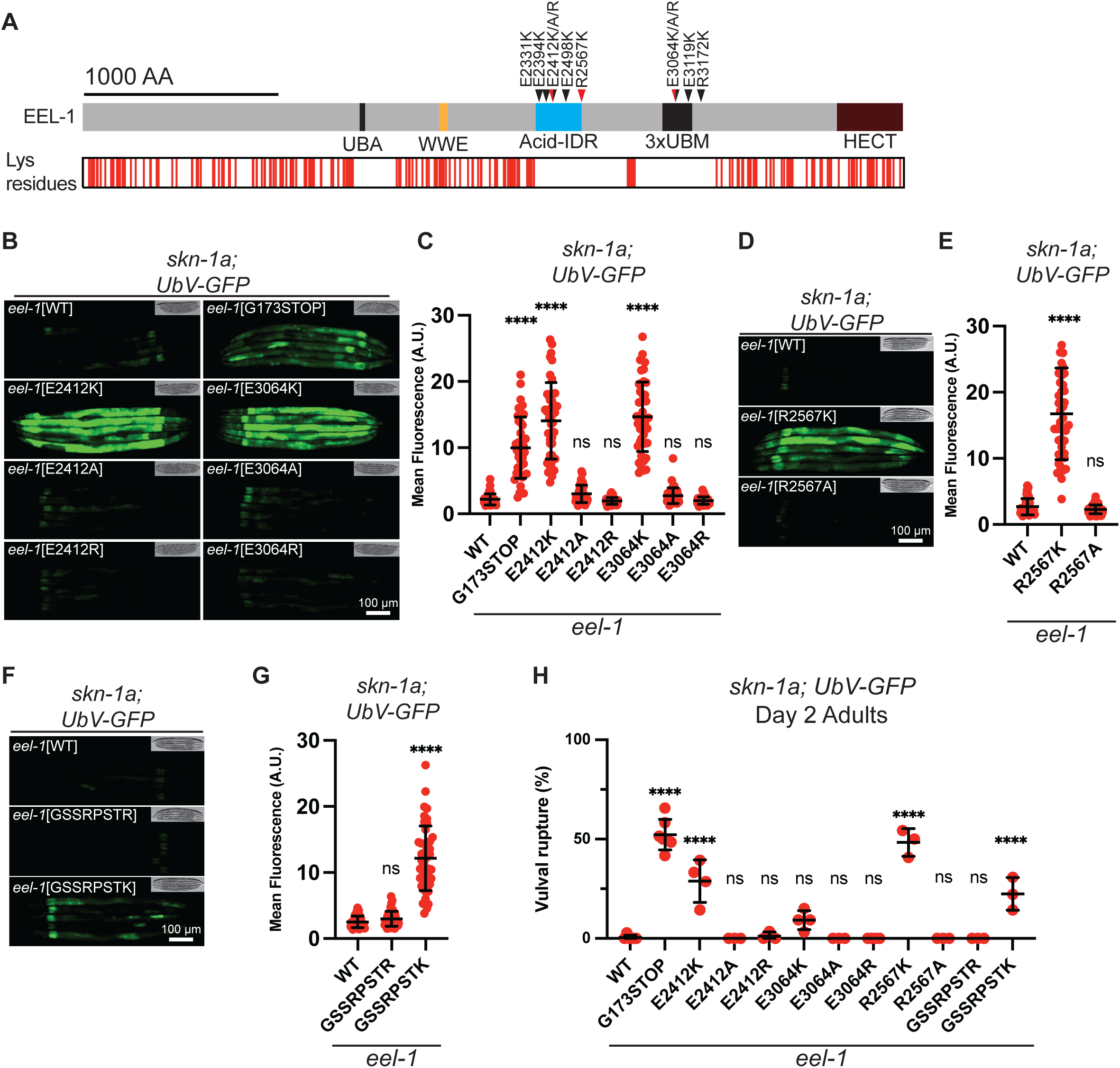
Lysine deficiency within EEL-1/HUWE1 lysine deserts is required for degradation of UbV-GFP and its disruption has varied effects on vulval integrity. A) Schematic showing the EEL-1 protein, showing locations of all lysine residues. The Acid-IDR and 3xUBM domains are located within extensive lysine-free regions. Locations of amino acid substitution mutations affecting these domains are indicated. Black arrows indicate mutations isolated via mutagenesis screening; red arrows indicate mutations generated by CRISPR/Cas9 gene editing. B) Fluorescence micrographs showing that introducing a lysine (K) at position E2412 or E3064 causes increased accumulation of UbV-GFP in the *skn-1a* mutant background, but that introducing alanine (A) or arginine (R) does not. Scale bar 100 μM. C) Quantification of UbV-GFP levels shown in (B). All animals are *skn-1a* mutant, the x-axis label indicates the *eel-1* genotype. Results are shown for n=45 animals. Error bars show mean ± SD. Ns p>0.5, ** p<0.01, **** p<0.0001, indicates P-value for comparison to the *eel-1*[WT] control, ordinary one-way ANOVA with Dunnett’s multiple comparisons test. D) Fluorescence micrographs showing that introducing a lysine (K) at position R2567 causes increased accumulation of UbV-GFP in the *skn-1a* mutant background, but that introducing alanine (A) does not. Scale bar 100 μM. E) Quantification of UbV-GFP levels shown in (D). All animals are *skn-1a* mutant, the x-axis label indicates the *eel-1* genotype. Results are shown for n=45 animals per genotype. Error bars show mean ± SD. Ns p>0.5, **** p<0.0001, indicates P-value for comparison to the *eel-1*[WT] control, ordinary one-way ANOVA with Dunnett’s multiple comparisons test. F) Fluorescence micrographs showing that insertion of an arbitrary lysine-free sequence, GSSRPSTR, at position 2410 does not cause increased accumulation of UbV-GFP in the *skn-1a* mutant background, but that introducing a similar lysine-containing sequence, GSSRPSTK, does. Scale bar 100 μM. G) Quantification of UbV-GFP levels shown in (F). All animals are *skn-1a* mutant, the x-axis label indicates the *eel-1* genotype. Results are shown for n=45 animals per genotype. Error bars show mean ± SD. Ns p>0.5, **** p<0.0001, indicates P-value for comparison to the *eel-1*[WT] control, ordinary one-way ANOVA with Dunnett’s multiple comparisons test. H) Quantification of vulval rupture phenotypes caused by various *eel-1* mutations affecting lysine deserts. All animals are *skn-1a* mutants and carry the UbV-GFP transgene, the x-axis label indicates the *eel-1* genotype. Results with the *eel-1*[*G173STOP*] allele are included for comparison. Results are shown for n=3-7 replicate experiments, 25-35 animals were assayed in each replicate experiment. Error bars show mean ± SD. Ns p>0.05, **** p<0.0001, indicates P-value for comparison to the *eel-1*[WT] control, ordinary one-way ANOVA with Šídák’s multiple comparisons test. Additional statistical analysis of these data is provided in Table S1.

We used CRISPR/Cas9 to recreate two of the mutations isolated in our screen; E2412K (a substitution within the Acid-IDR) and E3064K (within the 3xUBMs) (Fig 4A). As expected, both cause increased accumulation of UbV-GFP in the *skn-1a* mutant background (Fig 5B, C). Importantly, substitution of alanine (A) or arginine (R) at these positions has no effect (Fig 5B, C). These data suggest that disruption of EEL-1/HUWE1 function by these mutations is a specific consequence of introducing lysine to EEL-1’s lysine-free regions.

The lysine deficient regions of EEL-1 span hundreds of amino acids. We wondered whether our screen had pinpointed specific locations within the desert that are particularly sensitive to introduction of a lysine. To test this idea, we arbitrarily selected a basic residue within this region, R2567, which is at the periphery of the Acid-IDR domain but near the center of a lysine desert (Fig 5A). Strikingly, in a *skn-1a* mutant background, R2567K, but not R2567A, causes a strong enhancement of UbV-GFP accumulation (Fig 5D, E). The strong effect of introducing a lysine at a randomly selected location suggests that this lysine-deficient region of EEL-1/HUWE1 is intolerant of lysine at many or all positions, as implied by its deep evolutionary conservation.

Our data indicate that ectopic lysine residues in the Acid-IDR of EEL-1 are not tolerated, even though this domain can be deleted without disrupting function. This suggests that the ectopic lysine residues do not simply disrupt an Acid-IDR-dependent function.

Rather, EEL-1 function in general requires that this domain is lysine-free. To test this, we inserted a short lysine-free arbitrary sequence, GSSRPSTR, into the Acid-IDR after the 2410^th^ residue (*eel-1*[*PSTR^2410^*]), and a similar lysine-bearing variant insertion, GSSRPSTK (*eel-1*[*PSTK^2410^*]). Strikingly, only the lysine-bearing insertion causes a UbV-GFP accumulation in a *skn-1a* mutant background (Fig 5F, G). Thus, lysine residues, even if engineered to alter a novel sequence not normally present in the protein, still potently disrupt function. This confirms that the need for these regions to be lysine-free includes residues that have no direct relevance to the surrounding domain’s function(s). This suggests that conservation of the lysine-deficient domains is driven by strict selection to be lysine-free, which creates an additional sequence constraint on top of those imposed by the domains’ functions.

### Lysine desert mutations in eel-1 variably disrupt vulval integrity

To gauge the physiological impact of manipulations to the lysine desert of EEL-1, we measured their effect on vulval integrity in day 2 adults (Fig 5H). In the *skn-1a* genetic background, none of the non-lysine substitutions or insertions caused any increase in vulval rupture, consistent with little or no effect on *eel-1* function. In contrast, mutations that introduce an ectopic lysine cause vulval rupture in the *skn-1a* mutant background, albeit to very varying extents. The variant we engineered at a randomly selected position, R2567K, caused the strongest phenotype, equivalent to an *eel-1* null allele, but the amino acid substitutions isolated in our mutagenesis screen had less dramatic effects (Fig 5H, Table S1). The E2412K variant causes an intermediate rupture phenotype, whereas we did not observe elevated vulval rupture in E3064K mutants.

However, *eel-1*[*E3064K*] *skn-1a* double mutants did display markedly elevated vulval rupture by day 4 of adulthood (Fig S8). When analyzed as single mutants, neither *eel-1*[*E2412K*] or *eel-1*[*E3064K*] caused vulval rupture at day 4 of adulthood, a phenotype we had observed in the *eel-1* null, suggesting that these mutations do not fully inactivate EEL-1 (Fig 2C, Fig S8). Thus, the physiological impacts of lysine desert mutations are variable and dependent on the specific location at which ectopic lysine is introduced.

### Mutant phenotypes caused by an ectopic lysine residue are rescued by an engineered protein-protein interaction

Lysine deserts are often found in intrinsically disordered domains that are likely to be dynamically exposed to solvent and/or cellular interactors, properties that might predispose adventitious ubiquitination [81]. To investigate the relationship between disorder, solvent exposure, and lysine desert mutations’ functional impact, we inserted a proline-flanked ALFA tag (PSRLEEELRRRLTEP, a lysine-free engineered sequence with high helical propensity [87]) after the 2410^th^ residue of EEL-1 (*eel-1*[*ALFA^2410^*]). We generated a second edit in which the same insertion is preceded by a lysine residue (<u>K</u>PSRLEEELRRRLTEP; *eel-1*[*K-ALFA^2410^*]). We find that insertion of ALFA into the disordered Acid-IDR does not cause UbV-GFP accumulation in the *skn-1a* mutant background, unless accompanied by insertion of a lysine residue (Fig 6A, B).

**Figure 6.**
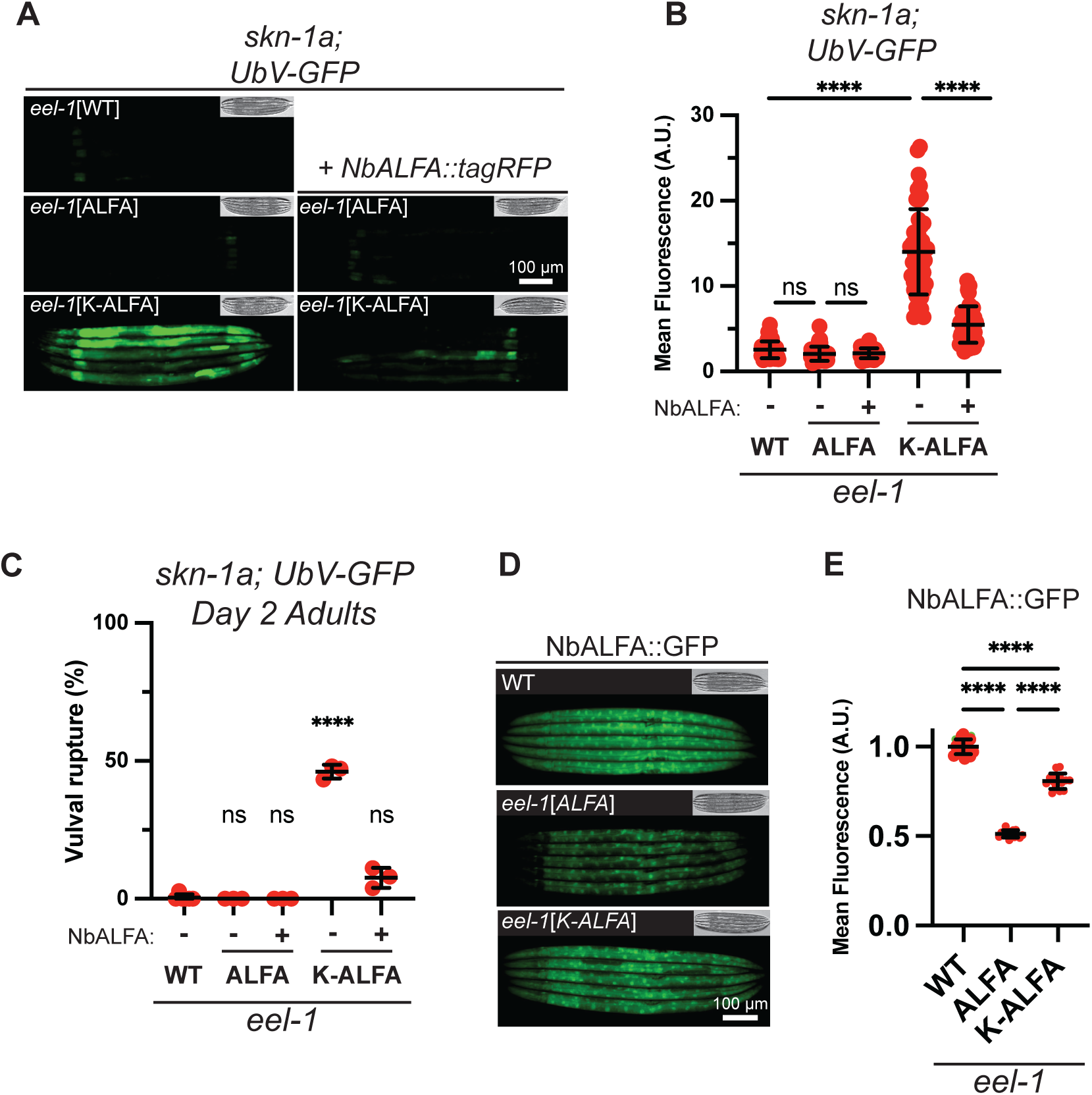
Engineered changes to the lysine desert of EEL-1/HUWE1 modulate UbV-GFP turnover and vulval integrity. A) Fluorescence micrographs showing that insertion of a proline-flanked ALFA tag (“ALFA”’; PSRLEEELRRRLTEP) at position 2410 does not cause increased accumulation of UbV-GFP in the *skn-1a* mutant background, but the same insertion accompanied by a flanking lysine residue (“K-ALFA”; KPSRLEEELRRRLTEP), does. The UbV-GFP accumulation caused by the K-ALFA insertion is substantially reduced by expression of NbALFA::tagRFP. Scale bar 100 μM. B) Quantification of UbV-GFP levels shown in (A). All animals are *skn-1a* mutant, the x-axis label indicates the *eel-1* genotype and the presence of NbALFA::tagRFP. Results are shown for n=45 animals per genotype. Error bars show mean ± SD. Ns p>0.5, **** p<0.0001, ordinary one-way ANOVA with Tukey’s multiple comparisons test. C) Quantification of vulval rupture phenotypes caused by engineered insertion of an ALFA tag in the EEL-1 lysine desert. All animals are *skn-1a* mutants and carry the UbV-GFP transgene, the x-axis label indicates the *eel-1* genotype and presence of NbALFA::tagRFP. Results are shown for n=3 replicate experiments, 25-35 animals were assayed in each replicate experiment. Error bars show mean ± SD. Ns p>0.05, **** p<0.0001, indicates comparison to the eel-1[WT] control, ordinary one-way ANOVA with Šídák’s multiple comparisons test. Additional statistical analysis of these data is provided in Table S1. D) Fluorescence micrographs showing that engineered insertion of an ALFA tag at position 2410 of *eel-1* leads to degradation of NbALFA::GFP, and that the extent of degradation is limited if the engineered insertion includes an adjacent lysine residue. Scale bar 100 μM. E) Quantification of NbALFA::GFP levels shown in (D). The x-axis label indicates the *eel-1* genotype. Results are shown for n=15 animals per genotype. Error bars show mean ± SD. **** p<0.0001, ordinary one-way ANOVA with Tukey’s multiple comparisons test

Consistently, in the *skn-1a* mutant background, *eel-1*[*K-ALFA^2410^*], but not *eel-1*[*ALFA^2410^*], causes vulval rupture (Fig 6C). In fact, the extent of vulval rupture caused by the K-ALFA insertion is higher than that caused by the non-helical GSSRPSTK insertion at the same location. Thus, a short helical insertion in the Acid-IDR does not impair function unless accompanied by an ectopic lysine residue. Moreover, the detrimental effect of an ectopic lysine at this location is not blunted, but rather exacerbated, if adjacent to an engineered helical insertion in the disordered Acid-IDR.

NbALFA, a single chain antibody, binds to the ALFA tag with very high affinity [87]. We wondered whether NbALFA binding would have any effect on the function of EEL-1[ALFA^2410^] or EEL-1[K-ALFA^2410^]. We hypothesized that tightly associating the Acid-IDR, which is disordered and lysine deficient, to a non-disordered and lysine-containing protein, could interfere with EEL-1[ALFA^2410^] function. Alternatively, for EEL-1[K-ALFA^2410^], placing a bulky binding partner adjacent to the ectopic lysine could modulate detrimental effects. We engineered animals to express NbALFA fused to tagRFP under the strong *rpl-28* promoter from a single-copy genomic insertion. Interestingly, NbALFA::tagRFP has no apparent effect on EEL-1[ALFA^2410^] function, as measured by accumulation of UbV-GFP or vulval rupture in a *skn-1a* mutant background (Fig 6A-C). In contrast, the defect in UbV-GFP turnover and the vulval rupture of *eel-1*[*K-ALFA^2410^*] animals is strongly suppressed by NbALFA::tagRFP (Fig 6A-C). This suggests that the detrimental effects of the ectopic lysine residue in EEL-1[K-ALFA^2410^] is dampened by NbALFA binding, likely by restricting the exposure of the ectopic lysine to the cellular milieu.

### Ectopic lysine impairs degradation of an engineered EEL-1 substrate

In EEL-1[ALFA^2410^]-expressing animals, binding of NbALFA fusion proteins to the ALFA tag would recruit them to the substrate recognition arena of EEL-1 and would likely cause their degradation. We examined the levels of an NbALFA::GFP fusion in *eel-1*[ALFA^2410^] animals and observed a significant reduction, suggesting that the engineered interaction between EEL-1 and NbALFA::GFP causes its ubiquitination and destruction. Interestingly, destabilization of NbALFA::GFP by EEL-1[K-ALFA^2410^] is diminished compared to EEL-1[ALFA^2410^], although both contain an intact ALFA sequence (Fig 6D, E). Thus, introduction of a lysine residue within the Acid-IDR domain interferes with degradation of an engineered substrate that binds adjacent to the ectopic lysine residue. These data suggest that the ectopic lysine residue reduces the ability of EEL-1 to ubiquitinate bound substrates and/or trigger their subsequent destruction.

### The effects of lysine desert mutations are not explained by reduced EEL-1 stability

Lysine deserts can safeguard against accidental ubiquitination leading to self-destruction of UPS components [81–86]. To address whether lysine amino acid substitutions affect the stability of EEL-1, we engineered amino acid substitutions at position E2412 into a strain in which endogenous EEL-1 is tagged with mNeonGreen (mNG::EEL-1) [88]. To control for non-specific effects of amino acid substitutions at position E2412 on EEL-1 stability, we generated E2412K, E2412A and E2412R variants. We did not detect any difference in mNG::EEL-1 levels or localization caused by any mutation at position E2412, including E2412K (Fig S9). This suggests a functional requirement for lysine deficiency at this position not related to stability.

## DISCUSSION

Proteins artificially fused to ubiquitin are subject to proteasomal degradation via the UFD pathway [53, 56, 57]. Analysis of UFD pathway reporters (such as UbV-GFP) indicate that the UPS is remarkably resilient to impairment of proteasome activity. In cancer cells, no accumulation of UbV-GFP is observed in cells even when approximately 80% of proteasomes are inactivated by a proteasome inhibitor drug [53]. Our analysis provides insight into an EEL-1/HUWE1-dependent mechanism that allows the UPS to tolerate partial proteasome inactivation. Further, our findings provide evidence that this mechanism safeguards UPS function to ensure normal animal development and physiology.

Physical association between HUWE1/Tom1 and UFD substrates has been demonstrated in human and yeast cells [58, 60]. In UFD and in protein quality control more generally, HUWE1/Tom1 appears to collaborate with other ubiquitin ligases to create ubiquitin chain(s) of appropriate length and/or topology to drive robust degradation [25, 58, 60, 89, 90]. Our genetic analysis indicates the participation of Ub-binding domains of EEL-1 in the degradation of UbV-GFP. This supports a model in which EEL-1/HUWE1/Tom1 directly binds engineered ubiquitin fusion proteins (or pre-ubiquitinated endogenous proteins) to facilitate further ubiquitination and promote their degradation. *In vitro*, ubiquitination of UFD substrates by Tom1 requires a second UFD ubiquitin ligase, Ufd4 [60]. HUWE1 preferentially modifies substrates already modified by long ubiquitin chains [25], and interactome studies suggest that HUWE1 binds ubiquitin chains of varied linkage topologies [27, 89]. Altogether, these observations suggest that EEL-1/HUWE1/Tom1 can function together with other E3 ligases to optimize protein degradation through amplification of ubiquitin chains and/or cooperative assembly of complex ubiquitin chain architectures.

Our findings, together with studies of human and fungal orthologs, suggest that EEL-1/HUWE1/Tom1 is particularly important for maintaining proteostasis when cellular capacity to maintain adequate protein turnover is challenged [24–26, 60]. Excess proteotoxic stress can stabilize ubiquitinated species [91], which could create the opportunity and/or need for EEL-1/HUWE1/Tom1 to catalyze further ubiquitination. In this way, EEL-1 could help ensure timely destruction of ubiquitinated proteins as cells adapt to environmental stressors or at critical moments in development. EEL-1/HUWE1 can promote degradation of exogenous proteins artificially fused to Ub, suggesting that any ubiquitinated protein could potentially be a target for further ubiquitination by EEL-1/HUWE1. However, precise substrate selection by EEL-1/HUWE1 *in vivo* could prioritize critical degradation pathways according to cellular needs. It will be of great interest to dissect the regulatory logic that dictates the specificity EEL-1/HUWE1 substrate selection in different physiological contexts.

Simultaneous inactivation of *eel-1* and *skn-1a* causes severe defects in UFD and the overall health of the animal that are not observed in either single mutant. Our data support a model in which SKN-1A and EEL-1 act in distinct pathways that each support optimal function of the UPS. The SKN-1A/Nrf1 pathway acts to maintain an adequate cellular pool of functional proteasomes via transcriptional control of proteasome biogenesis. In contrast, the functional deficit in EEL-1 mutants is not accompanied by a reduction in proteasome levels, and our degron-GFP degradation assay suggests that proteasomes are still functionally competent. These data further support a crucial role for EEL-1/HUWE1-dependent ubiquitination in the regulatory network that adapts the UPS to fulfill cellular needs.

UPS dysfunction – that is, failure or misregulation of the UPS regulatory network – is a hallmark of aging and a cause of age-associated loss of proteostasis [5, 6]. Age-dependent vulval integrity defects are a marker of loss of tissue homeostasis during aging in *C. elegans* [92]. Our observations show that whereas *eel-1* and *skn-1a* single mutants show elevated vulval rupture in an age-dependent manner [16], double mutants show a severe defect in vulval integrity as young adults. Catastrophic enhancement of this age-dependent phenotype in double mutants reinforces the view that EEL-1/HUWE1 and SKN-1A/Nrf1 act in a complementary manner to maintain proteostasis and animal health during aging. It will be important to identify the proteasomal substrate(s) that underlie the synergistic effects of EEL-1 and SKN-1A inactivation and define their roles in development, physiology, and aging.

UPS dysfunction is linked to abnormalities in human neuronal development, function, and adult-onset neurodegenerative diseases. Mutations affecting the Nrf1 pathway, HUWE1, or proteasomal components themselves cause an overlapping spectrum of neurodevelopmental symptoms [50–52, 77, 93–102]. Both the SKN-1A/Nrf1 pathway and EEL-1/HUWE1 are protective against aggregation of proteins implicated in late-onset proteinopathies such as Alzheimer’s Disease and Huntington’s Disease [14, 16, 25, 101, 103–106]. Pharmacological interventions that increase HUWE1 levels or activity may have therapeutic potential in neurological conditions. Alternatively, manipulating HUWE1 substrate specificity could represent a general avenue for therapeutic manipulation of the UPS.

Protein domains entirely deficient in lysine residues are a conserved feature of eukaryotic proteomes and are frequently found in proteins that participate in UPS function [81]. However, lysine deserts’ relevance to animal development and physiology have been unclear, and their specific functional roles are not well understood. Our genetic analysis demonstrates that the absence of lysine residues from the lysine desert of EEL-1 is important for this Ub-dependent Ub ligase to optimize UPS function.

Interestingly, a lysine-introducing mutation in *eel-1* (E2475K, within the Acid-IDR lysine desert) alters neuronal architecture, supporting relevance to neuronal development and function [41].

We have not directly measured whether lysine residues placed within the lysine desert of EEL-1 are subject to ubiquitination. Such ubiquitination events might be difficult to detect, since EEL-1/HUWE1/Tom1 readily undergoes autoubiquitination [46–48, 107]. Despite this limitation, our findings are most consistent with a model in which lysine residues introduced to the lysine deserts of EEL-1 undergo ubiquitination, as observed for lysine deserts in other UPS components [81–86]. Ubiquitination within lysine deserts may cause degradation [81–86], but analysis of the E2412K mutation revealed no evidence of reduced EEL-1 levels, indicating some other effect on function. One possibility is that ubiquitination of EEL-1 within the lysine desert occludes functionally important interactions with substrates or other regulators. However, our data argue against this possibility, as the E2412K mutation is not phenocopied by either (1) deletion of the surrounding lysine-deficient domain, or (2) engineering this part of EEL-1 to interact with an exogenously expressed NbALFA fusion protein. Another possibility is that ubiquitination of the ectopic lysine residues competes with substrate ubiquitination. The beneficial effect of NbALFA fusion proteins, which would be expected to limit access for ubiquitination of the ectopic lysine in EEL-1[K-ALFA^2410^], is consistent with this model. Altogether, our findings support the model proposed by Kampmeyer and colleagues, in which UPS components contain lysine deserts to allow them to function in proximity to ubiquitin ligases without competing with genuine substrates for ubiquitination [81].

We have generated and characterized several mutations that introduce a lysine residue to lysine deserts of EEL-1 at different locations. All cause strong accumulation of UbV-GFP, but their impact on physiology is variable. This variable phenotypic impact hints that ectopic lysine residues at different positions do not simply diminish EEL-1 activity to differing extents but rather have separation-of-function effects. Possibly, differently positioned ectopic lysine residues compete with different subsets of genuine substrates for ubiquitination. Our mutagenesis screen selected mutations that cause defective UbV-GFP turnover but do not severely impact development (mutants that were inviable or very difficult to culture were discarded). This appears to have biased towards uncovering a set of *eel-1* variants that very strongly disrupt UbV-GFP turnover while sparing other degradation pathways that are important for normal physiology.

Interestingly, drug-like small molecules that are ubiquitinated by HUWE1 can interfere with endogenous substrates’ ubiquitination and degradation [108]. Conceivably, targeting such molecules to different locations in the HUWE1 substrate-binding arena could have distinct effects, analogous to differently positioned ectopic lysine residues.

Many proteins containing lysine deserts are highly conserved throughout eukaryotic evolution [81]. Lysine deserts are also present in bacterial species that regulate protein turnover via pupylation, suggesting they were an early molecular adaptation to protein degradation via lysine-linked protein conjugation [83]. Inspecting the sequences of EEL-1/HUWE1/Tom1 orthologs reveals deep conservation of domain architecture and sequence compositional biases. This conservation extends to these domains’ structural arrangement, as revealed by experimentally determined structures of human, nematode, and fungal orthologs [46–48]. This implies that the common ancestor of EEL-1/HUWE1/Tom1 contained a HECT domain accompanied by lysine-deficient Ub-binding and Acid-IDR domains approximately 1.5 billion years ago. The ancient origin of EEL-1/HUWE1/Tom1 argues that its present-day functional versatility exists because it has repeatedly been coopted into degradation pathways that must function optimally to support the complexities of eukaryotic biology.

## METHODS

### C. elegans strains and maintenance

*C. elegans* were maintained on standard nematode growth media (NGM) at 20°C and fed *E. coli* OP50 unless otherwise noted. A list of strains used in this study is provided in Table S2. Some strains were provided by the CGC, which is funded by NIH Office of Research Infrastructure Programs (P40 OD010440). Ethane-methyl-sulfonate (EMS) mutagenesis was carried out by treating ∼10,000 L4 animals with 47 mM EMS for 4 hours at 20°C. After mutagenesis, large populations of synchronized F2 animals were screened and individuals with increased accumulation of UbV-GFP were isolated. This screen was carried out using *skn-1a*(*mg570*); *mgIs77*[*UbV-GFP*] and in *png-1*(*ok1654*); *mgIs77*[*UbV-GFP*] animals.

### Whole genome sequencing and analysis

Genomic DNA was prepared using the Gentra Purgene Tissue kit (Qiagen, #158689). Genomic DNA libraries were prepared using the NEBNext genomic DNA library construction kit (New England Biolabs, #E6040). Libraries were sequenced using an Illumina HiSeq instrument. Deep sequencing reads were analyzed using a custom Galaxy workflow adapted from CloudMap [109]. Candidate causative alleles were identified by the isolation of multiple independent mutations affecting the same gene.

### Growth rate assay

Embryos were isolated by bleaching of gravid adult animals and then hatched overnight in M9 at 20°C while rotating. The resulting population of synchronized L1 larvae were plated to fresh NGM plates seeded with OP50 and incubated at the desired temperature. After 48-72 hours, plates were imaged using a Leica M165FC equipped with a 910 Leica K5 sCMOS camera and using LAS X software. Animal length was measured using the segmented line tool in ImageJ.

### Vulval rupture assay

To measure adult rupture, 30-40 L4 animals were selected at random from mixed stage cultures and transferred to a fresh plate. The animals were monitored for rupture (indicated by intestinal or germline material outside the vulva) after 48 hours (day 2 adults), and again after a further 48 hours (day 4 adults). Throughout the rupture assay, adults were transferred to fresh plates every 2-3 days to separate them from progeny.

To image vulval rupture in day 1 adults, L4 animals were selected at random from mixed stage cultures and transferred to a fresh plate. After 24 hours, the animals were imaged using a Leica M165FC equipped with a 910 Leica K5 sCMOS camera and using LAS X software.

### Plasmid constructs and transgenesis

All cloning was carried out using the NEBuilder HiFi DNA Assembly kit (New England Biolabs #E2621). The sequence encoding NbALFA was codon optimized for expression in *C. elegans* using the transgenebuilder tool (https://www.wormbuilder.org/transgenebuilder/) and synthesized to include a single intron (from the *rps-0* gene) and a C-terminal in-frame linker sequence (STSGGSGGTGGSS). Gene synthesis was carried out by Twist Biosciences. Gibson assembly was used to generate plasmids in which NbALFA fusion proteins are expressed under control of the ubiquitous *rpl-28* promoter (605 bp immediately upstream of the *rpl-28* ORF) and are followed by the *tbb-2* 3’UTR (376 bp immediately downstream of the *tbb-2* stop codon). These constructs were inserted to the pNL43 backbone, to allow generation of transgenics by miniMos [15, 110]. pNL559 (Rpl-28p::NbALFA::tagRFP::tbb-2) and pNL549 (Rpl-28p::NbALFA::GFP::tbb-2), were used to generate transgenic (miniMos insertion) animals harboring single-copy transgenes driving ubiquitous expression of NbALFA::tagRFP and NbALFA:tagRFP respectively [110].

The *pbs-5_p_::gfp* reporter was generated by placing the promoter of CEOP1752 (850 bp upstream of the first open reading frame of the operon, *K05C4.2* (*pbs-5* is the second gene in the operon), upstream of a fragment containing the GFP coding sequence and *tbb-2* 3’UTR (376 bp immediately downstream of the *tbb-2* stop codon), and inserted to the pBlueScript backbone, to generate pNL136 (*pbs-5_p_::GFP*). High-copy-number arrays containing pNL136 were integrated to chromosome II via CRISPR/Cas9 [111, 112].

### Genome Modification by CRISPR/Cas9

CRISPR/Cas9 genome editing was carried out through microinjection of guide RNA/Cas9 ribonucleoprotein (RNP) complexes (IDT #1081058, #1072532, and custom crRNA) along with single-stranded oligonucleotides (IDT) as homology-directed repair (HDR) templates [113, 114]. Sequences of all guide RNAs and HDR oligos are provided in Table S3. All CRISPR edits were identified by diagnostic PCR and confirmed by sequencing.

### Microscopy

Brightfield and fluorescence images of whole animals (UbV-GFP, *pbs-5_p_::gfp*, *rpn-9::mScarlet*, degron-GFP, NbALFA::GFP) were collected on a Leica M165FC equipped with a 910 Leica K5 sCMOS camera and using LAS X software. Higher magnification images (mNG::EEL-1) were collected on a Ziess AxioImager Z1 microscope equipped with an Axiocam 705 CMOS camera and using ZEN software. For fluorescence imaging, worms were immobilized using sodium azide and mounted on 2% agarose pads. All images were processed and analyzed using ImageJ/Fiji software. Images shown within the same figure panel were collected using the same exposure time and were then processed identically. To quantify accumulation of UbV-GFP, the mean pixel intensity in the anterior intestinal cells was measured. To quantify *pbs-5_p_::GFP* reporter expression, *rpn-9::mScarlet* levels, degron-GFP levels, and NbALFA::GFP levels, the mean pixel intensity was measured using a polygon selection around each animal. To quantify mNG::EEL-1 levels, the mean pixel intensity was measured in 2-3 circular sections corresponding to the nuclei of the proximal 2-3 oocytes.

### Bortezomib treatment

NGM plates containing the desired concentration of bortezomib (LC Laboratories, #B1408) were prepared by directly applying bortezomib solution to NGM plates seeded with OP50 and leaving the plates to dry for 1-2 hours. Control plates supplemented with DMSO were prepared in parallel. L4 stage animals to were shifted to bortezomib or control DMSO-supplemented plates and imaged after 24 hours.

### Auxin-induced degradation

NGM plates containing 50 mM auxin (α-Napthaleneacetic acid, Phytotech Labs #N610) were prepared by directly adding auxin solution to NGM plates seeded with OP50 and leaving plates to dry for 1-2 hours. Control (non-auxin-exposed) L4 animals were imaged immediately prior to beginning the auxin treatment. L4 stage animals were shifted to auxin-supplemented plates and imaged after 1, 2 and 4 hours.

### Protein sequence analysis

EEL-1/HUWE1/Tom1 orthologs were identified in the PANTHER database subfamily: E3 UBIQUITIN-PROTEIN LIGASE HUWE1 (PTHR11254:SF67) [115]. Protein sequences of EEL-1 and orthologs were retrieved from Uniprot [116]. Multiple sequence alignment was carried out using Clustal Omega with default settings [117]. AIUPred was used to predict intrinsically disordered regions [118]. Sequence composition was analyzed in excel. Structure prediction was carried out using Alpha Fold 3 and structure predictions were visualized using UCSF ChimeraX [119, 120].

### Statistical analysis

Statistical tests were carried out using GraphPad Prism. All biological replicates were performed using independent populations of animals. All numerical data used in graphs and statistical tests are provided in Supplementary Data File S1.

## ACKNOWLEDGEMENTS

We thank Wormbase for *C. elegans* genome data and curation. Some strains were provided by the CGC, which is funded by NIH Office of Research Infrastructure Programs (P40 OD010440). The *png-1*(*ok1654*) mutant was generated by the was provided by the *C. elegans* Gene Knockout Project at the Oklahoma Medical Research Foundation, which was part of the International *C. elegans* Gene Knockout Consortium [121]. We thank Julián Cerón for sharing *rpn-9*(*cer203*[*rpn-9::mScarlet*]).

## FUNDING

This work was supported by the National Institutes of Health (NIH) National Institute of General Medical Sciences (grant R35GM142728 to NL). KSY was supported by the Biological Mechanisms of Healthy Aging (BMHA) training grant (T32AG066574).

---

**Figure S1.**
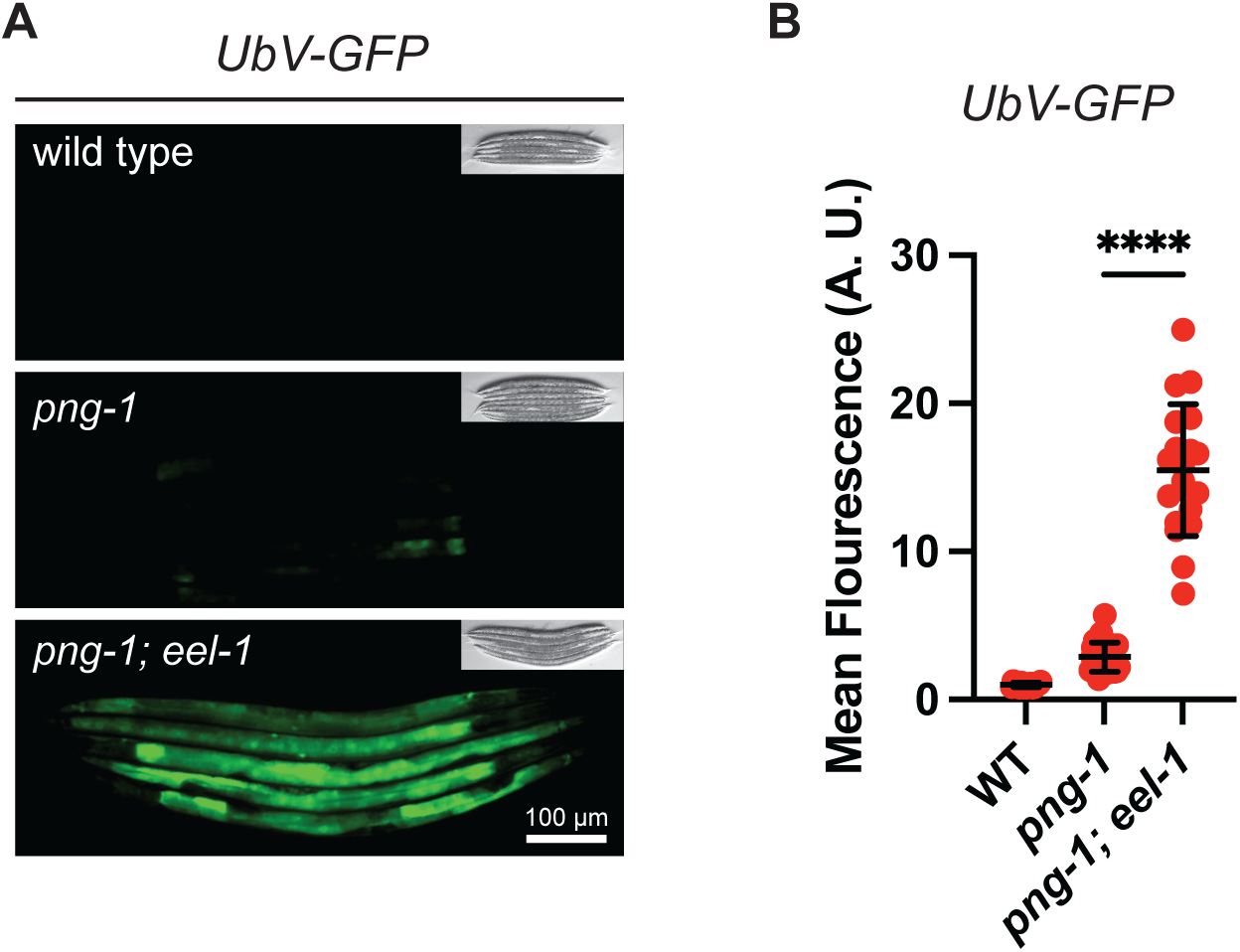
Dramatic accumulation of UbV-GFP in *png-1; eel-1* double mutants. A) Fluorescence micrographs showing accumulation of UbV-GFP in *png-1*; *eel-1* double mutants. All images are of L4 stage animals. Scale bar 100 μM. B) Quantification of UbV-GFP levels shown in (A). UbV-GFP is dramatically increased in *png-1*; *eel-1* double mutants. Error bars show mean ± SD. n>15 animals per genotype. **** p<0.0001, Welch’s t test.

**Figure S2.**
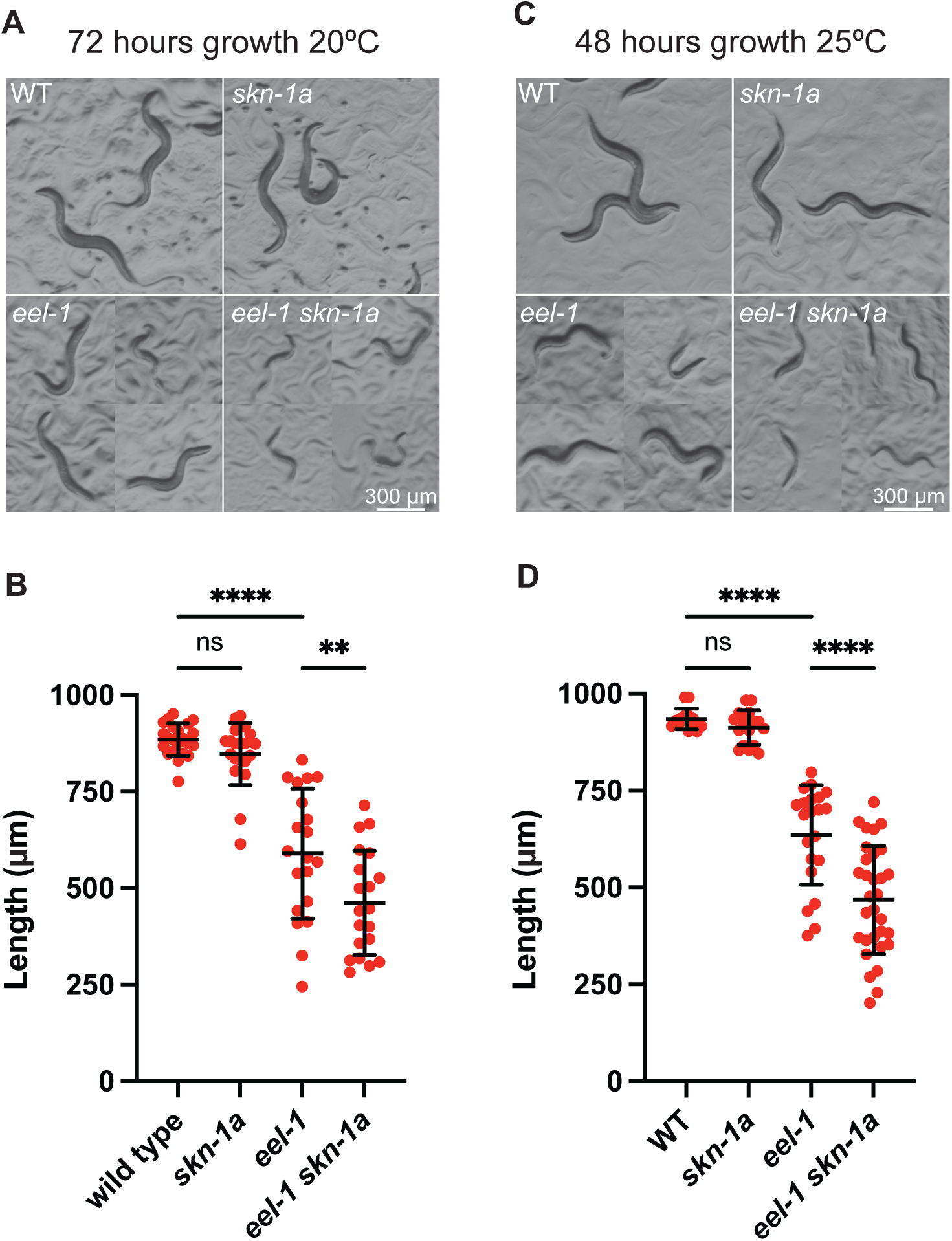
Growth defects of *eel-1* mutants are enhanced by *skn-1a* inactivation. A) Images showing the effect of *skn-1a* and *eel-1* mutations on developmental progression. Animals were synchronized at the L1 stage by hatching in buffer overnight and then cultured in the presence of OP50 bacteria for 72 hours at 20°C. Scale bar 300μM. B) Quantification of animal length from (A). The developmental progression of *skn-1a* mutants is identical to the wild type. The developmental progression of *eel-1* mutants is delayed, and this effect is exacerbated in *eel-1 skn-1a* double mutants. Error bars show mean ± SD. n=20 animals per genotype. Ns p>0.05; **** p<0.0001, ordinary one-way ANOVA with Šídák’s multiple comparisons test. C) Images showing the effect of *skn-1a* and *eel-1* mutations on developmental progression. Animals were synchronized at the L1 stage by hatching in buffer overnight and then cultured in the presence of OP50 bacteria for 48 hours at 25°C. Scale bar 300 μM. D) Quantification of animal length from (C). The developmental progression of *skn-1a* mutants is identical to the wild type. The developmental progression of *eel-1* mutants is delayed, and this effect is exacerbated in *eel-1 skn-1a* double mutants. Error bar shows mean ± SD. n>13 animals per genotype. Ns p>0.05; **** p<0.0001, ordinary one-way ANOVA with Šídák’s multiple comparisons test.

**Figure S3.**
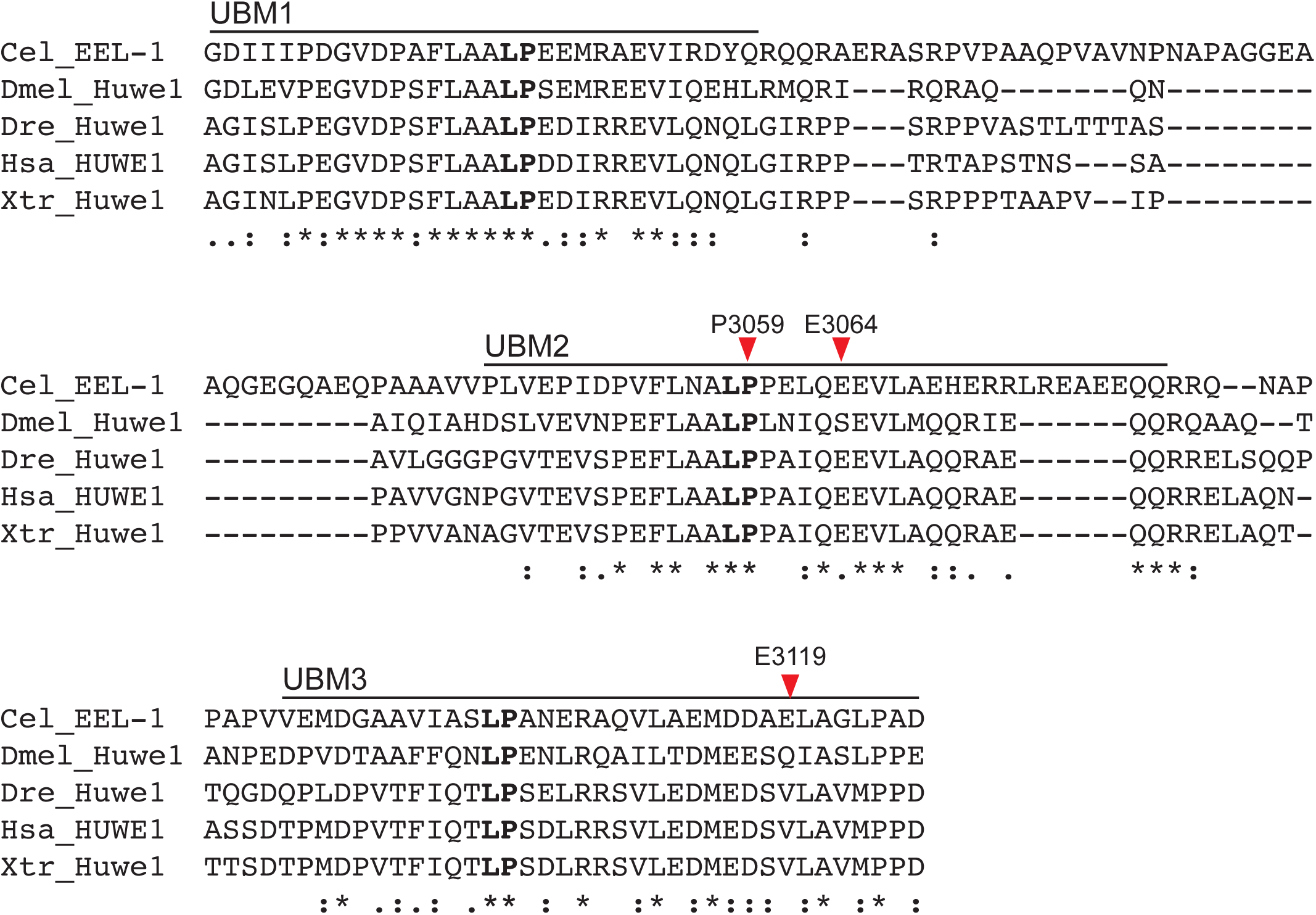
Sequence conservation of the EEL-1/HUWE1/Tom1 3xUBM domain. Multiple sequence alignment of EEL-1/HUWE1/Tom1 3xUBM domain. The invariant LP motif found in each UBM is in bold. Locations of amino acid substitution mutations analyzed in this study are indicated by red arrows.

**Figure S4.**
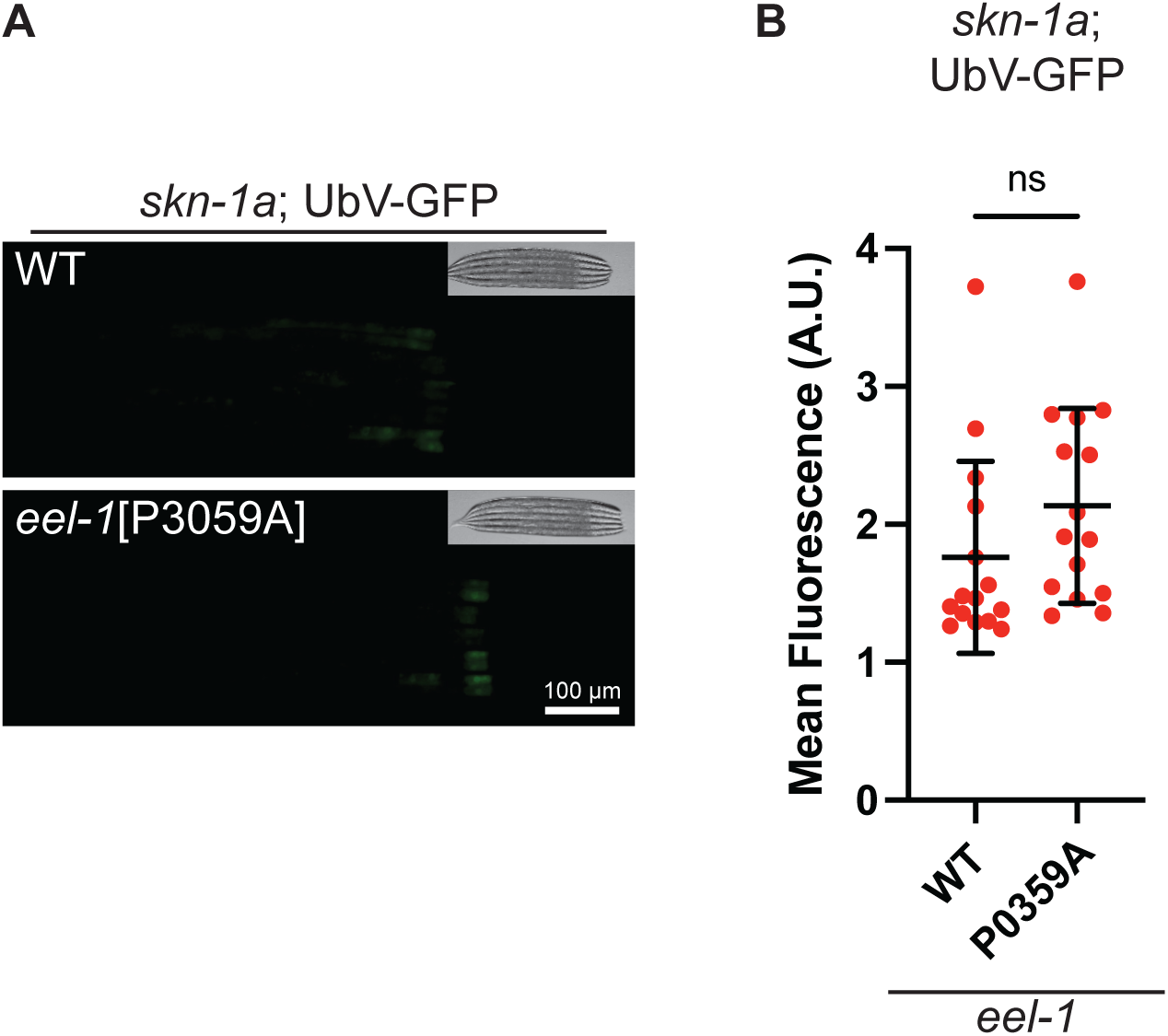
Eel-1[P3059A] mutation does not disrupt UbV-GFP degradation in the *skn-1a* mutant background. A) Fluorescence micrographs showing accumulation of UbV-GFP is not increased by the *eel-1*[*P3059A*] mutation in the *skn-1a* mutant background. Scale bar 100 μM. B) Quantification of UbV-GFP levels shown in (A). All animals are *skn-1a* mutant, the x-axis label indicates the *eel-1* genotype. Error bars show mean ± SD. Results are shown for n=15 animals. Ns p>0.05, Welch’s t test.

**Figure S5.**
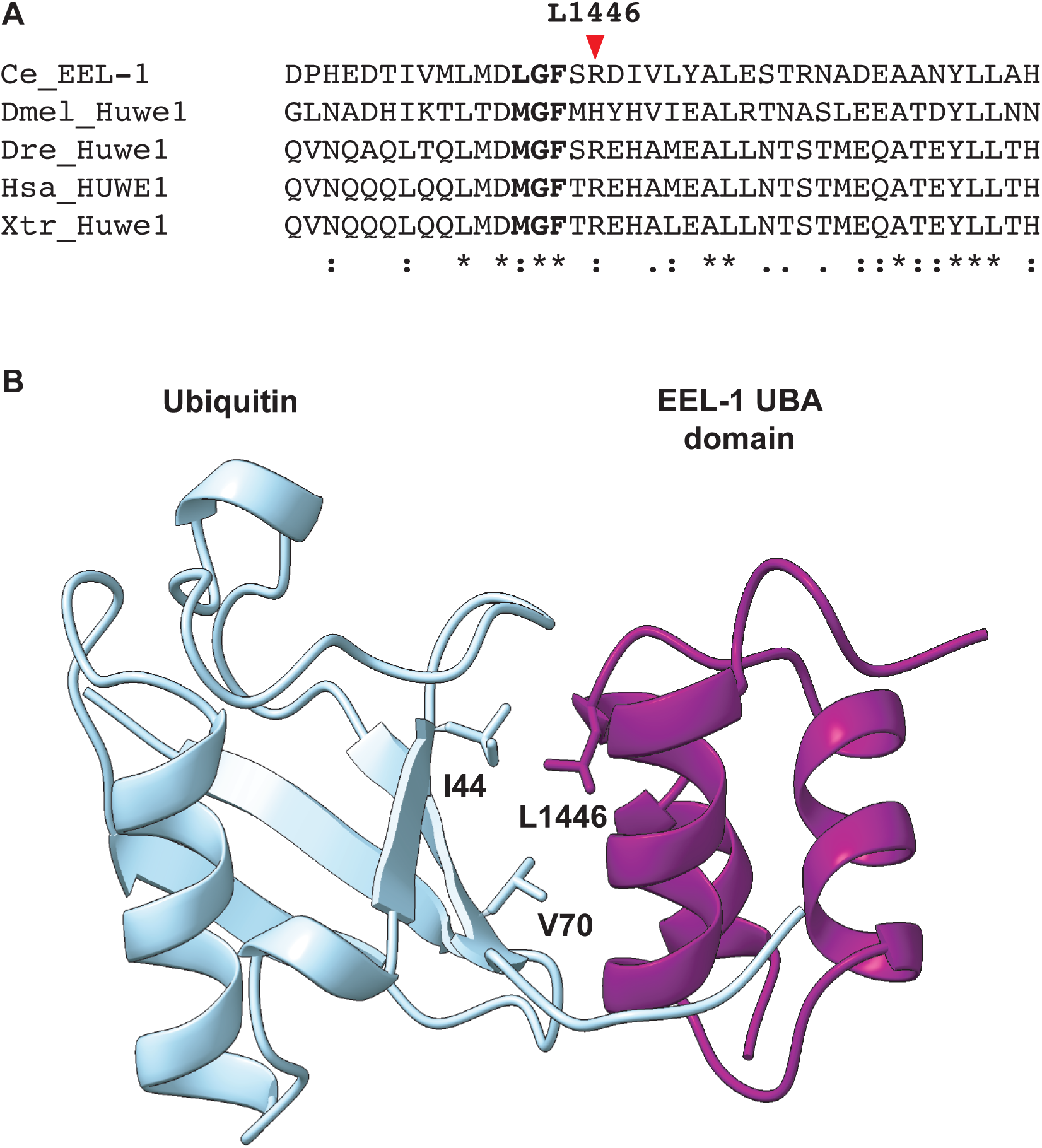
Sequence conservation of the EEL-1/HUWE1/Tom1 UBA domain. A) Multiple sequence alignment of EEL-1/HUWE1/Tom1 UBA domain. The conserved L/MGF motif is in bold. L1446 is indicated by a red arrow. B) Alpha Fold 3 model showing predicted interaction between ubiquitin and the EEL-1 UBA domain. L1446 and hydrophobic residues of ubiquitin involved in the ubiquitin:UBA interaction are labelled.

**Figure S6.**
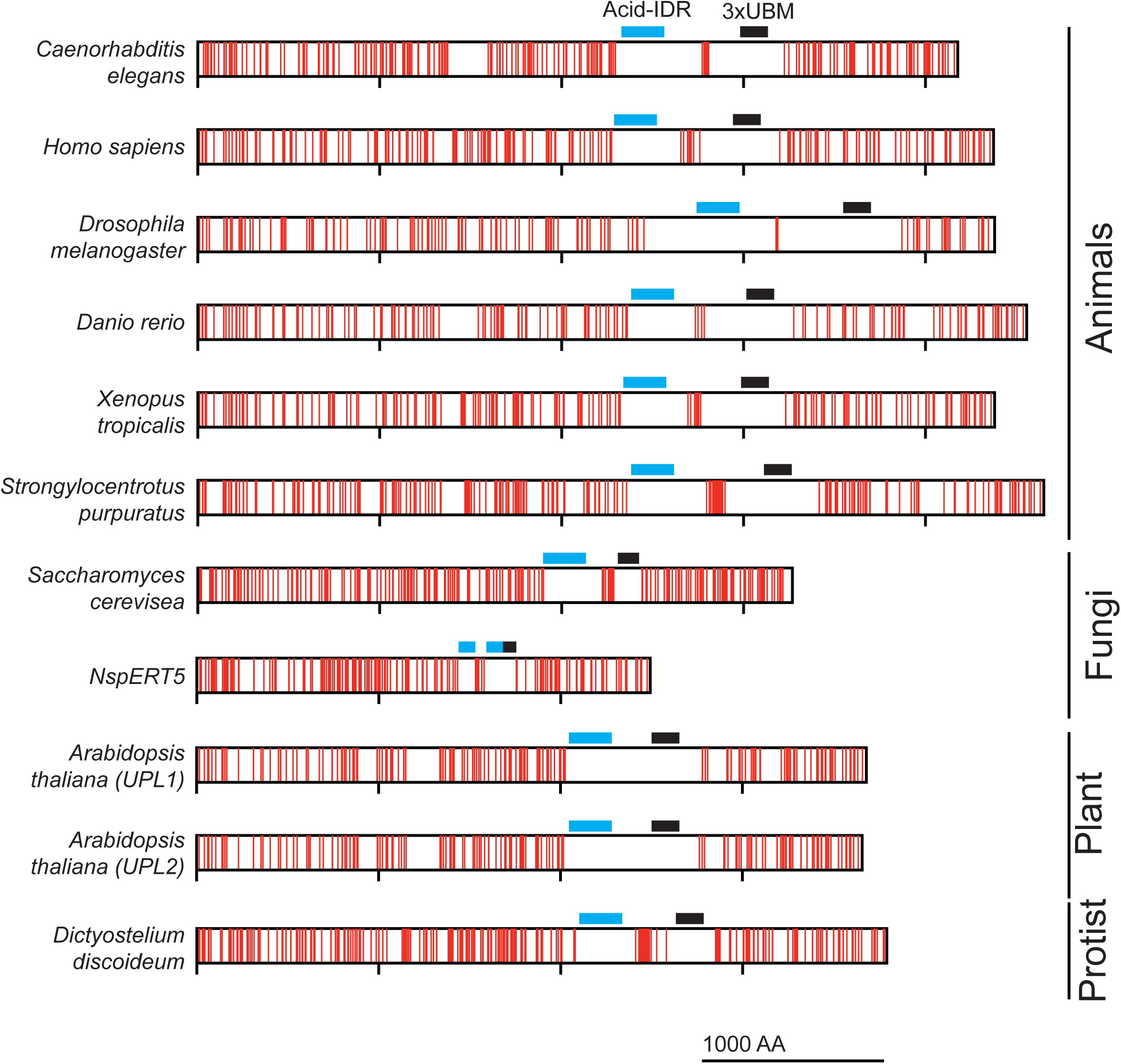
Amino acid composition of EEL-1 and orthologs showing conserved lysine deficient regions. For each protein, red lines indicate locations of lysine residues. The locations of the Acid-IDR domain (blue) and 3xUBM domain (black) of each ortholog is indicated. The location of the Acid-IDR was inferred by analysis of each ortholog’s sequence composition (see Fig S7).

**Figure S7.**
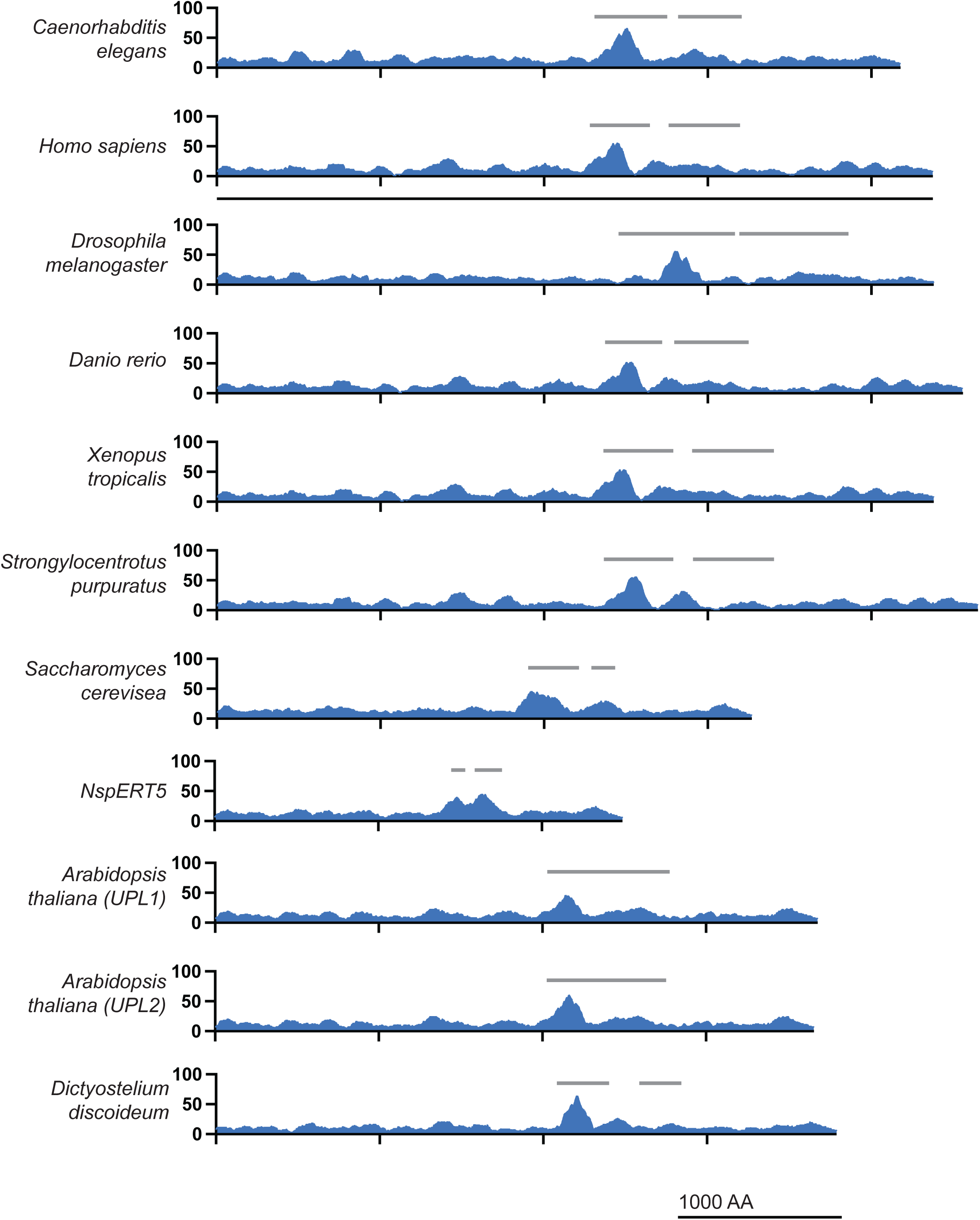
Amino acid composition of EEL-1 and orthologs showing conserved regions enriched in acidic amino acids. For each protein, the percentage acidic (glutamate or aspartate) residues within a moving 100 amino acid window is plotted. Grey lines indicate lysine-deficient regions of each ortholog (see Fig S6).

**Figure S8.**
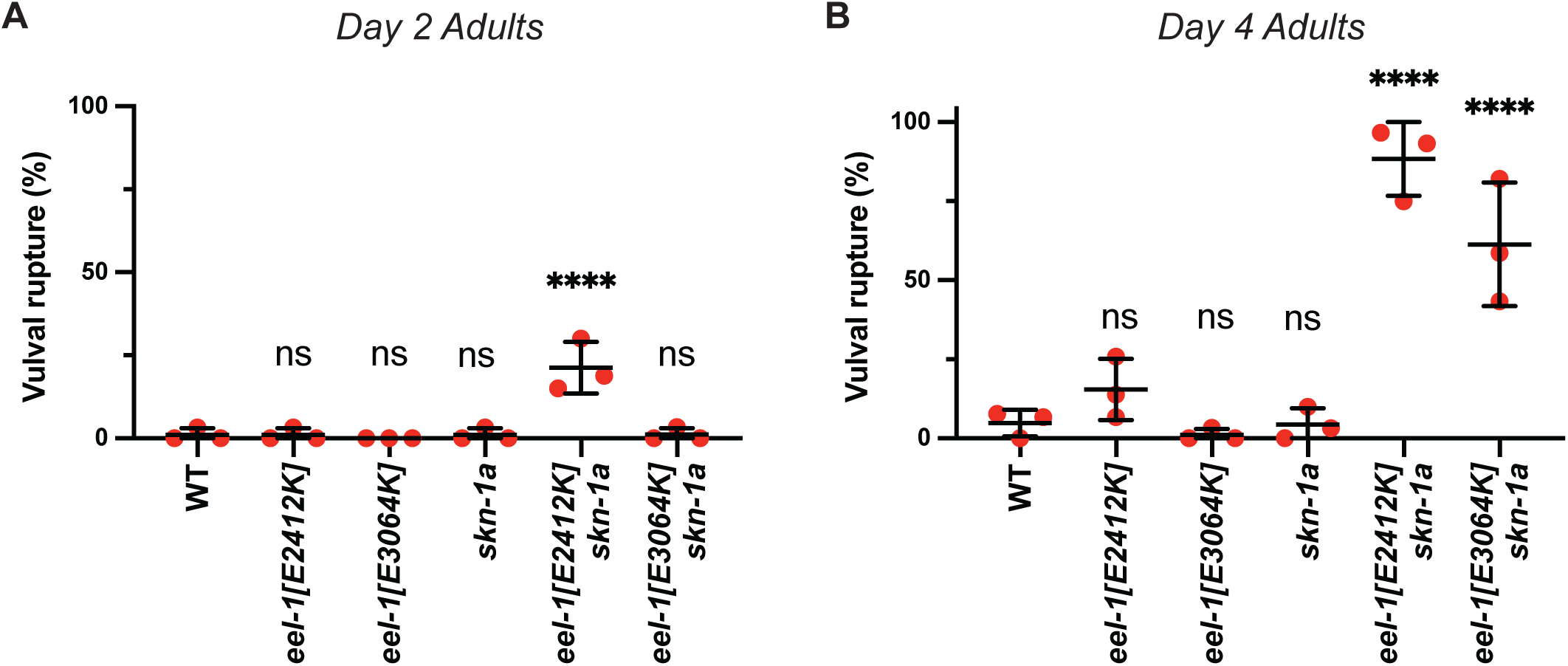
Vulval rupture of *eel-1*[*E2412K*] and *eel-1*[*E3064K*] mutants. A) Quantification of vulval rupture in day 2 adults. Results are shown for n=3 replicate experiments, 25-35 animals were assayed in each replicate. Error bars show mean ± SD. Ns p>0.05, **** p<0.0001, indicates P-value compared to the wild type (WT) control, ordinary one-way ANOVA with Šídák’s multiple comparisons test. B) Quantification of vulval rupture in day 4 adults. Results are shown for n=3 replicate experiments, 25-35 animals were assayed in each replicate. Error bars show mean ± SD. Ns p>0.05, **** p<0.0001, indicates P-value compared to the wild type (WT) control, ordinary one-way ANOVA with Šídák’s multiple comparisons test.

**Figure S9.**
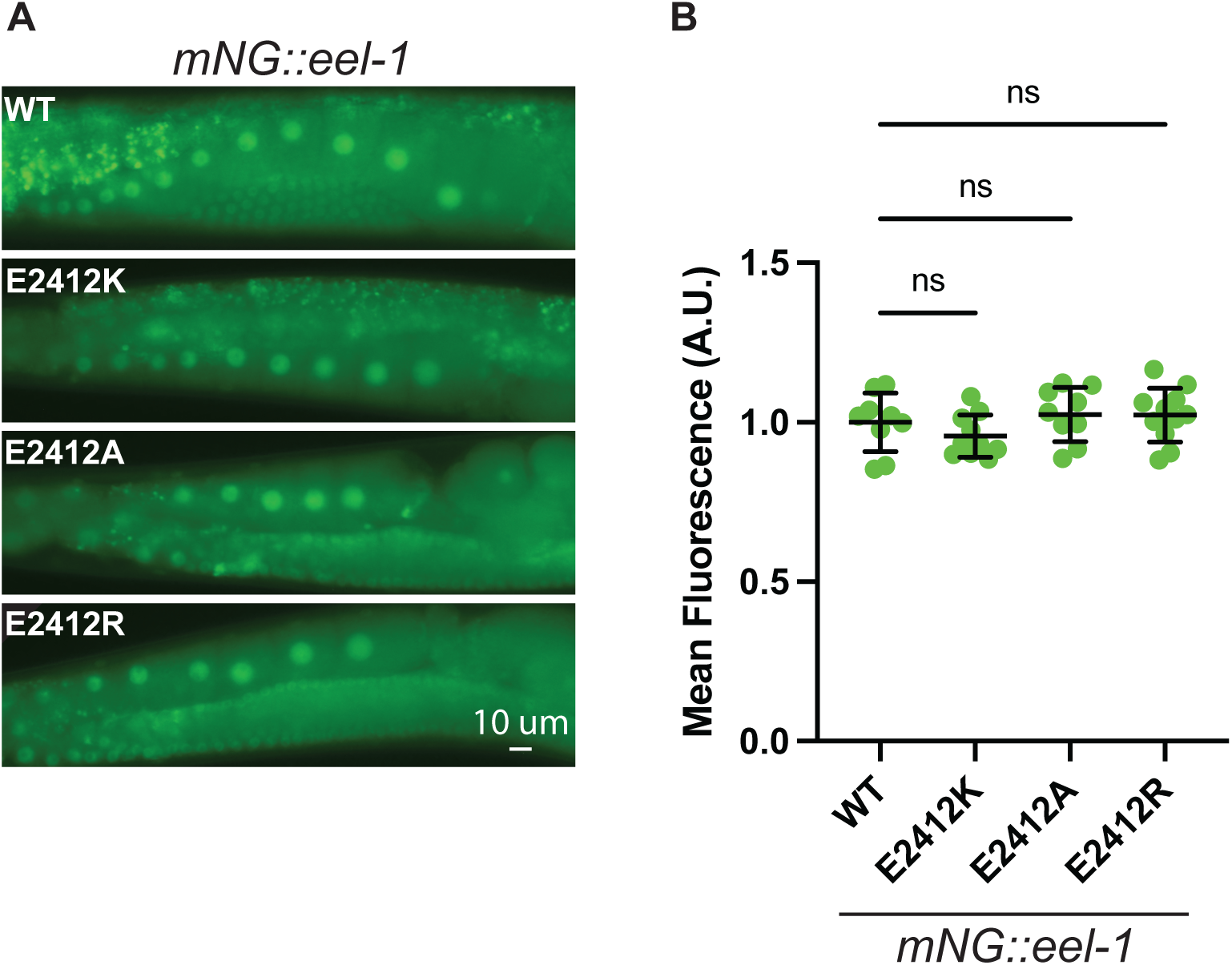
Amino acid substitutions at E2412 do not alter localization or expression of mNG::EEL-1. A) Fluorescence micrographs showing endogenously tagged mNG::EEL-1. Images show germline of adult animals, which show prominent expression of mNG-tagged EEL-1, which is predominantly localized to the nucleus. The expression and localization of mNG::EEL-1 is not altered by amino acid substitutions at E2412. Scale bar 10 μM. B) Quantification of mNG::EEL-1 levels shown in (A). The prominent mNG::EEL-1 fluorescence present in oocytes was measured. Error bars show mean ± SD. Results are shown for n=9-11 nuclei for each genotype. Ns p>0.05, indicates P-value compared to the mNG::EEL-1[WT] control, ordinary one-way ANOVA with Dunnett’s multiple comparisons test.

## Notes

### Competing Interest Statement

The authors have declared no competing interest.

